# Taste triggers a homeostatic temperature control in hungry flies

**DOI:** 10.1101/2022.09.30.510382

**Authors:** Yujiro Umezaki, Sergio Hidalgo, Erika Nguyen, Tiffany Nguyen, Jay Suh, Sheena S. Uchino, Joanna C. Chiu, Fumika N. Hamada

## Abstract

Hungry animals consistently show a desire to obtain food. Even a brief sensory detection of food can trigger bursts of physiological and behavioral changes. However, the underlying mechanisms by which the sensation of food triggers the acute behavioral response remain elusive. We have previously shown in *Drosophila* that hunger drives a preference for low temperature. Because *Drosophila* is a small ectotherm, a preference for low temperature implies a low body temperature and a low metabolic rate. Here, we show that taste sensing triggers a switch from a low to a high temperature preference in hungry flies. We show that taste stimulation by artificial sweeteners or optogenetics triggers an acute warm preference, but is not sufficient to reach the fed state. Instead, nutrient intake is required to reach the fed state. The data suggest that starvation recovery is controlled by two components: taste-evoked and nutrient-induced warm preferences, and that taste and nutrient quality play distinct roles in starvation recovery. Animals are motivated to eat based on time of day or hunger. We found that clock genes and hunger signals profoundly control the taste-evoked warm preferences. Thus, our data suggest that the taste-evoked response is one of the critical layers of regulatory mechanisms representing internal energy homeostasis and metabolism.

## INTRODUCTION

Animals are constantly sensing environmental stimuli and changing their behavior or physiology based on their internal state ^1–7^. Hungry animals are strongly attracted to food. Immediately after seeing, smelling, or chewing food, even without absorbing nutrients, a burst of physiological changes is suddenly initiated in the body. These responses are known in mammals as the cephalic phase response (CPR) ^8^. For example, a flood of saliva and gastrointestinal secretions prepares hungry animals to digest food ^9–13^. Starvation results in lower body temperatures, and chewing food triggers a rapid increase in heat production, demonstrating CPR in thermogenesis ^14–16^. However, the underlying mechanisms of how the sensation of food without nutrients triggers the behavioral response remain unclear.

To address this question, we used a relatively simple and versatile model organism, *Drosophila melanogaster*. Flies exhibit robust temperature preference behavior ^17,18^. Due to the low mass of small ectotherms, the source of temperature comes from the environment. Therefore, their body temperatures are close to the ambient temperature ^19,20^. For temperature regulation, animals are not simply passive receivers of ambient temperature. Instead, they actively choose an ambient temperature based on their internal state. For example, we have shown that preferred temperature increases during the daytime and decreases during the night time, exhibiting circadian rhythms of temperature preference (temperature preference rhythms: TPR) ^21^. Because their surrounding temperature is very close to their body temperature, TPR leads to body temperature rhythms (BTR) that is very similar to mammalian BTR ^22,23^. Another example is the starvation. Hunger stress forces flies to change their behavior and physiological response ^24,25^. We previously showed that the hungry flies prefer a lower temperature ^26^. The flies in a lower environmental temperature has been shown to have a lower metabolic rate, and the flies in a higher environmental temperature has been shown to have a higher metabolic rate ^26–28^. Therefore, hungry flies choose a lower temperature and therefore, their metabolic rate is lower. Similarly, in mammals, starvation causes a lower body temperature, hypothermia ^6^. In mammals, body temperature is controlled by the balance between heat loss and heat production. The starved mammals have been shown a lower heat production ^5–7^. Therefore, both flies and mammals, the starvation causes a low body temperature.

The flies exhibit robust feeding behaviors ^29,30^ and molecular and neural mechanisms of taste are well documented ^2,31–36^. Therefore, we focused on taste and temperature regulation and asked how the taste cue triggers a robust behavioral recovery of temperature preference in starving flies. We show in hungry flies that taste without nutrients induces a switch from a low to a high temperature preference. While taste leads to a warmer temperature preference, nutrient intake causes the flies to prefer an even warmer temperature. This nutrient-induced warm preference results in a complete recovery from starvation. Thus, taste-evoked warm preference is different from nutrient-induced warm preference and potentially similar physiology as CPR. Therefore, when animals emerge from starvation, they use a two-step approach to recovery: taste-evoked and nutrient-induced warm preference. While a rapid component is elicited by food taste alone, a slower component requires nutrient intake.

Animals are motivated to eat based on their internal state, such as time of day or degree of hunger. The circadian clock drives daily feeding rhythms ^37–39^ and anticipates meal timing. Daily feeding timing influences energy homeostasis and metabolism ^40,41^. Circadian clocks control feeding behavior in part via orexigenic peptidergic/hormonal regulation such as neuropeptide Y (NPY) and agouti-related peptide (AgRP) neurons, which are critical for regulating feeding and metabolism ^39,42^. Feeding-fasting cycles modulate peripheral organs in liver, gut, pancreas, and so on ^43^, suggesting that circadian clocks, peptidergic/hormonal signals, and peripheral organs are organically coordinated to enable animals maintain their internal states constantly. We found that clock genes and hunger signals are strongly required for taste-evoked warm preference. The data suggest that taste-evoked response is an indispensable physiological response that represents internal state. Taken together, our data shed new light on the role of the taste-evoked response and highlight a crucial aspect of our understanding of feeding state and energy homeostasis.

## RESULTS

### Food detection triggers a warm preference

To investigate how food detection influences temperature preference behavior in *Drosophila* ^17,18^, the *white^1118^* (*w^1118^*) control flies were fed fly food containing carbohydrate, protein, and fat sources (Fig. 1A and S1) and tested in temperature preference behavioral assays (Fig. S1) ^26,44^. The flies are released into a chamber set to a temperature range of 16-34°C and subsequently accumulate at their preferred temperature (Tp) at 25.2±0.2 °C within 30 minutes (min) (Fig. 1B: fed, white bar). On the other hand, when *w^1118^* flies were starved overnight with water only, they preferred 21.7±0.3 °C (Fig. 1A and 1B: overnight starvation (STV), gray bar). Thus, starvation leads to a lower Tp. As we have previously reported, starvation strongly influences temperature preference ^26^.

**Figure 1.**
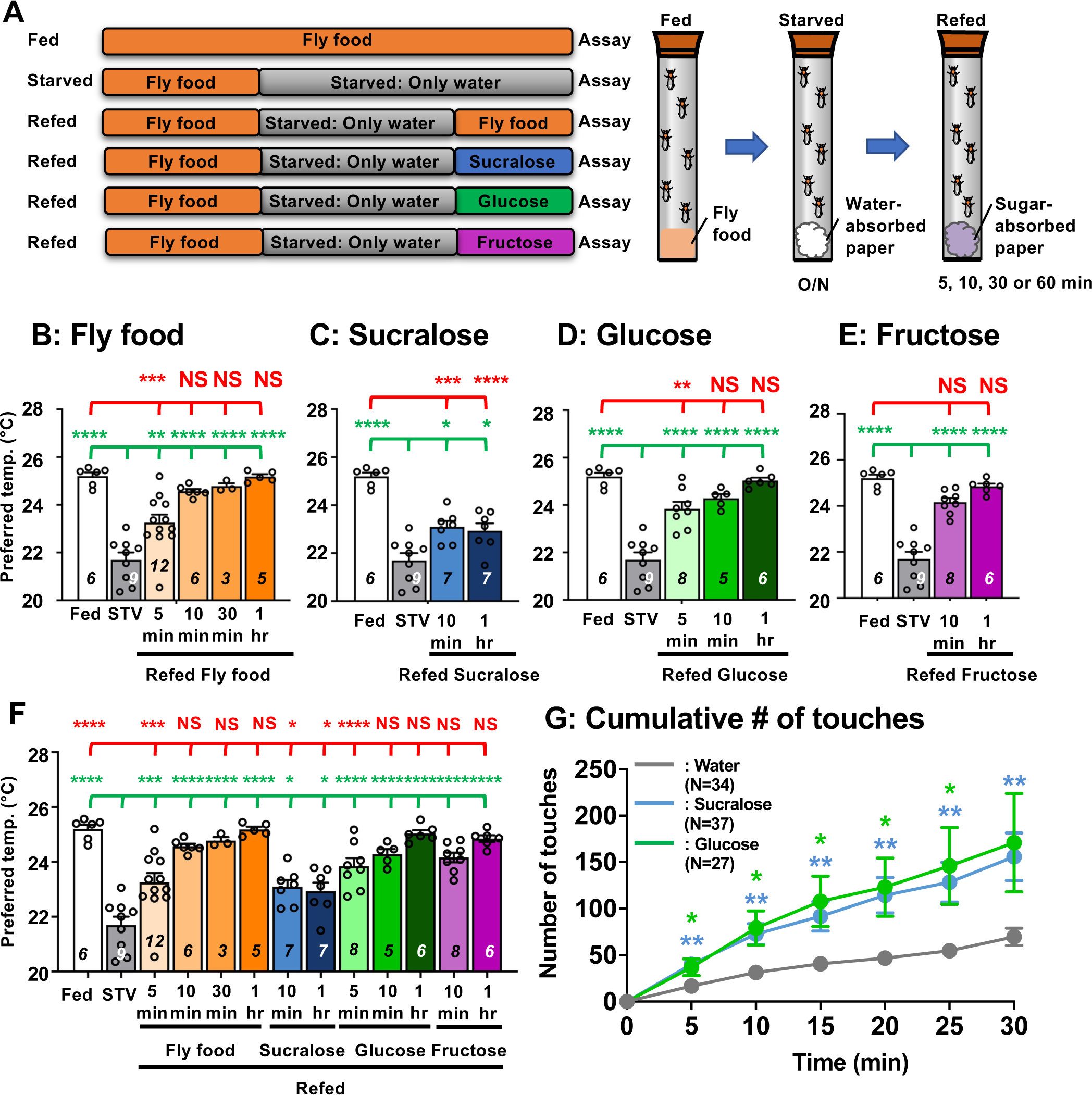
Hungry flies switch from cold to warm preference upon food detection. **(A)** A schematic diagram of the experimental conditions. Flies were reared on fly food. Emerging flies were kept on fly food for one or two days (orange) and then starved with water-absorbing paper. Starvation was applied for 1-3 overnights (ON). The duration of starvation was chosen when the flies showed a statistically significant difference in Tp between fed and starved conditions (see “Starvation conditions” in Materials and methods for details). Starved flies were refed with either fly food (orange bar), 2 mL sucralose (blue), glucose solution (green), or fructose solution (purple) absorbed paper and then examined for temperature preference (Tp) behavioral assays. Details are provided in the experimental procedures. **(B-E)** Comparison of preferred temperature (Tp) of *white^1118^* (*w^1118^*) control flies between fed (white bar), starved (STV; gray bar), and refed (orange, blue, or green bar) states. Starvation was applied for 1ON. Starved flies were refed with fly food (orange bar) for 5, 10, 30, or 60 min (1 hr) (B), 2.8 mM sucralose solution (blue bar) for 10 min or 1 hr (C), 2.8 mM (equivalent to 5%) glucose solution (green bar) for 5 min, 10 min, or 1 hr (D), or 2.8mM fructose solution (purple bar) for 10 min or 1 hr (E). Behavioral experiments were performed at the specific time points, Zeitgeber time (ZT) 4-7. ZT0 and ZT12 are light on and light off, respectively. Dots on each bar indicate individual Tp in the assays. Numbers in italics indicate the number of trials. The Shapiro-Wilk test was performed to test for normality. One-way ANOVA was performed to compare Tp between each refeeding condition. Red or green stars indicate Tukey’s post hoc test comparing differences between experimental and fed or starved conditions, respectively. Data are presented as mean Tp with SEM. *p < 0.05. **p < 0.01. ***p < 0.001. ****p < 0.0001. NS indicates no significance. **(F)** Comparison of Tp among the following conditions in *w^1118^* (control) flies: fed (white bar), starved overnight (O/N) (water or STV; gray bar), refed with fly food (fly food; orange bar), sucralose (blue bars), glucose (green bars), or fructose (purple bars). The same data are used in Fig. 1B-E. A distribution of temperature preference in each condition is shown in Fig. S2A. The Shapiro-Wilk test was performed to test for normality. One-way ANOVA was performed to compare Tp between each refeeding condition. Red (or green) stars indicate Tukey’s post hoc test comparing differences between experimental and fed (or starved) conditions, respectively. *p < 0.05. **p < 0.01. ***p < 0.001. ****p < 0.0001. NS indicates not significant. **(G)** Feeding assay: The number of touches to water (gray), 2.8 mM (equivalent to 5%) glucose (green), or 2.8 mM sucralose in each solution (blue) was examined using *w^1118^* flies starved for 24 hrs. Water, glucose and sucralose were tested individually in the separate experiments. A cumulative number of touches to water or sugar solution for 0-30 min was plotted. Two-way ANOVA was used for multiple comparisons. Blue and green stars show Fisher’s LSD post hoc test comparing sucralose (blue stars) or glucose (green stars) solution feeding to water drinking. All data shown are means with SEM. *p < 0.05. **p < 0.01. ***p < 0.001. ****p < 0.0001.

To examine how starved flies recover from lower Tp, they were offered fly food for 5 min, 10 min, 30 min, and 1 hr (Fig. 1A). Immediately after the flies were refed, the temperature preference behavior assay was examined. The assay takes 30 min from the time the flies are placed in the apparatus until the final choice is made. For example, in the case of 5 min refed flies, it took a total of 35 min from the start of refeeding to the end of the assay. After 10 min, 30 min, or 1 hr of fly food refeeding, starved flies preferred a temperature similar to that of fed flies (Fig. 1B, 1F, and S2: statistics shown as red stars, Table S1), suggesting a full recovery from starvation. On the other hand, refeeding after 5 min resulted in a warmer temperature than the starved flies. Nevertheless, Tp did not reach that of the fed flies (Fig. 1B, 1F, and S2). Therefore, refeeding the flies for 5 min resulted in a partial recovery from the starved state (Fig. 1B, 1F, and S2, statistics shown as green stars, Table S1). Thus, our data suggest that food intake triggers a warm preference in starved flies.

### Sucralose refeeding promotes a warm preference

While only 5 min of refeeding fly food caused hungry flies to prefer a slightly warmer temperature, 10 min of refeeding caused hungry flies prefer a similarly warmer temperature as the fed flies (Fig. 1B, 1F, and S2). Therefore, we hypothesized that food-sensing cues might be important for the warm preference. Sucralose is an artificial sweetener that activates sweet taste neurons in *Drosophila* ^45^ and modulates taste behaviors such as the proboscis extension reflex ^46–49^. Importantly, sucralose is a non-metabolizable sugar and has no calories. Therefore, to investigate how food-sensing cues are involved in warm preference, we examined how sucralose refeeding changes the temperature preference of starved flies. After starved flies were refed sucralose for 10 min or 1 hr (Fig. 1A), they preferred a warmer temperature; however, Tp was halfway between Tps of fed and starved flies (Fig. 1C, F and S2: blue, Table S1). Thus, sucralose ingestion induces a warm preference but shows a partial recovery of Tp from the starved state.

While refeeding the flies with food resulted in a full recovery of Tp from the starved state, refeeding them with sucralose resulted in only a partial recovery of Tp. Therefore, starved flies may use both taste cues and nutrients to fully recover Tp from the starved state. To evaluate this possibility, we used glucose, which contains both sweetness (gustatory cues) and nutrients (i.e., metabolizable sugars) (Fig. 1A), and tested glucose refeeding for 5 min, 10 min, and 1 hr. We found that 10 min or 1 hr glucose refeeding resulted in full recovery of Tp from the starved state (Fig. 1D, 1F and S2: statistics shown in red NS, Table S1) and was significantly different from starved flies (Fig. 1D, 1F and S2: statistics shown in green stars, Table S1). Thus, our data showed that sucralose refeeding induced partial recovery and glucose refeeding induced full recovery from the starved state.

It is still possible that the starved flies consumed glucose faster than sucralose during the first 10 min, which could result in a different warming preference. To rule out this possibility, we examined how often starved flies touched glucose, sucralose, or water during the 30 min using the Fly Liquid-food Interaction Counter (FLIC) system ^50^. The FLIC system assays allow us to monitor how much interaction between the fly and the liquid food reflects feeding episodes. We found that starved flies touched glucose and sucralose food at similar frequencies and more frequently than water during the 30 min test period (Fig. 1G, Table S1). The data suggest that flies are likely to feed on glucose and sucralose at similar rates. Therefore, we concluded that the differential effect of sucralose and glucose refeeding on temperature preference was not due to differences in feeding rate.

Furthermore, to confirm that sweet taste is more important than sugar structure, we instead used fructose, another simple sugar that contains sweetness and nutrients. Glucose and fructose are monosaccharides and a member of hexose and pentose, respectively. We tested fructose refeeding for 10 min and 1 hr. We found that 10 min of fructose refeeding resulted in full recovery of Tp from the starved state (Fig. 1E, 1F and S2: statistics shown in red NS, Table S1), and 10 min and 1 hr fructose refeeding were significantly different from starved flies (Fig. 1E, 1F and S2: statistics shown in green stars, Table S1). The data suggest that sweet taste is more important than the structures of the sugar compounds.

### Activation of sweet taste neurons leads to warm preference

To determine how taste elicits a warm preference, we focused on the sweet gustatory receptors (Grs), which detect sweet taste. We used sweet Gr mutants and asked whether sweet Grs are involved in taste-evoked warm preference. Two different sweet Gr mutants, *Gr5a^−/−^; Gr64a^−/−^*and *Gr5a^−/−^;;Gr61a^−/−^, Gr64a-f^−/−^*, are known to reduce sugar sensitivity compared to the control ^47,48,51,52^. We found that sweet Gr mutant flies exhibited a normal starvation response in which the Tp of starved flies was lower than that of fed flies (Fig. 2A and 2B, white and gray bars, statistics shown as green stars, Table S1). However, starved sweet Gr mutant flies did not increase Tp after 10 min sucralose refeeding (Fig. 2A and 2B, blue bars, statistics shown as green and red stars, Table S1). These data suggest that sweet Grs are involved in taste-evoked warm preference.

**Figure 2.**
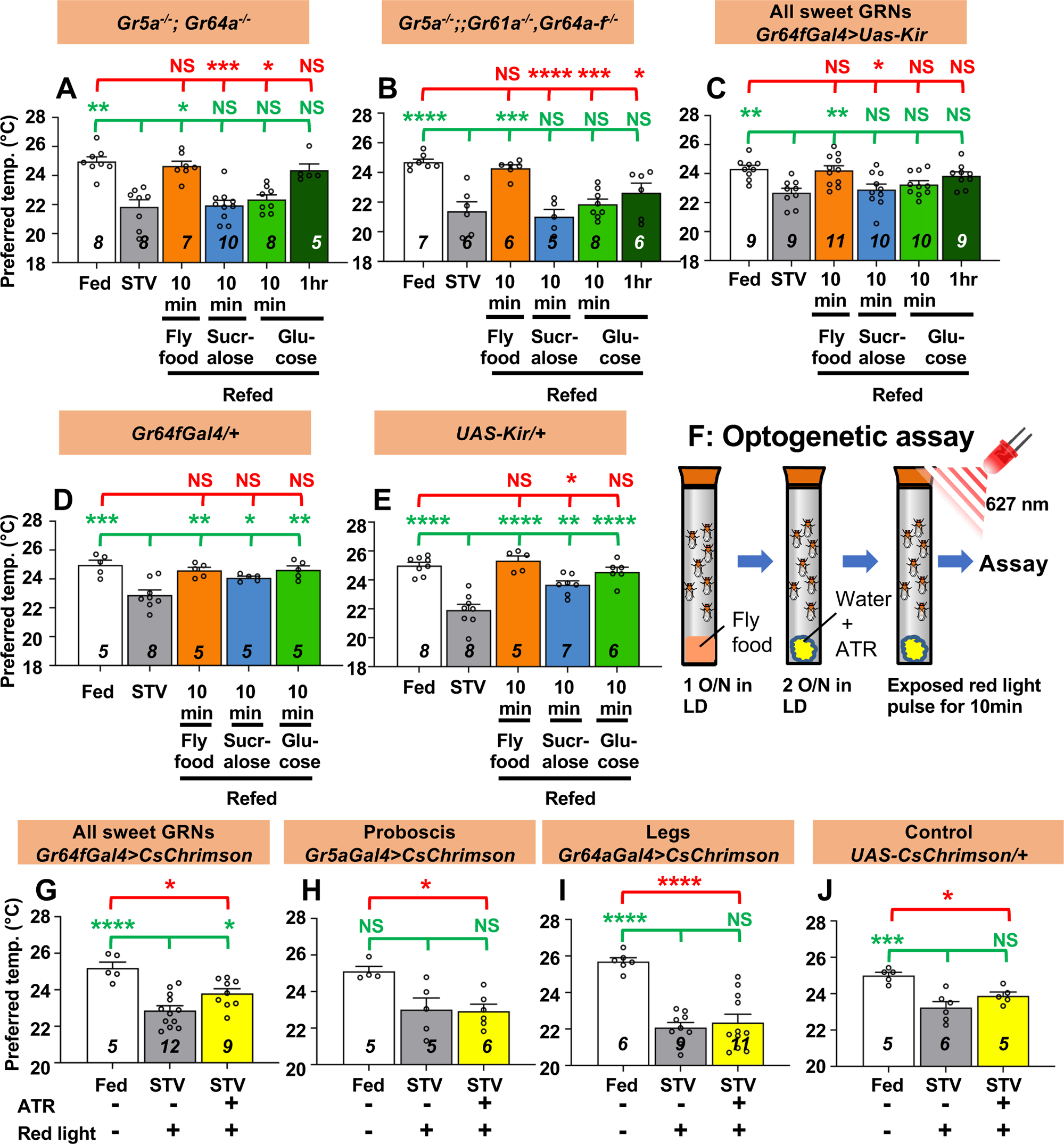
Gustatory neurons are essential for taste-evoked warm preference. **(A-E)** Comparison of preferred temperature (Tp) of flies between fed (white bar), starved (STV; gray bar), and refed (orange, blue, green, or dark green bar) states. Flies were starved for 2 overnights (ON) except for *Gr5a^−/−^;;Gr61a^−/−^, Gr64a-f^−/−^* (1.5ON). Starved flies were refed with fly food for 10 min (fly food; orange bars), sucralose for 10 min (blue bars), or glucose for 10 min (green bars), or 1 hr (dark green bars). **(F)** Schematic of the optogenetic activation assay. **(G-J)** Comparison of Tp of flies between fed (white bar), starved (STV; gray bar), and starved with all-trans-retinal (ATR; yellow bars), which is the chromophore required for CsChrimson activation. Gustatory neurons in starved flies were excited by red light pulses (flashing on and off at 10 Hz) for 10 min. Starvation was performed for 2ON. Averaged fly distributions of flies in the temperature gradient for *Gr64fGal4>CsChrimson* were shown in Fig. S2B. Behavioral experiments were performed on ZT4-7. Dots on each bar indicate individual Tp in assays. Numbers in italics indicate the number of trials. The Shapiro-Wilk test was used to test for normality. One-way ANOVA was used for statistical analysis. Red or green stars indicate Tukey’s post hoc test compared between each experiment and the fed (red) or starved (green) condition. All data presented are means with SEM. *p < 0.05. **p < 0.01. ***p < 0.001. ****p < 0.0001. NS indicates not significant.

Sweet Grs are expressed in the sweet Gr-expressing neurons (GRNs) located in the proboscis and forelegs ^52,53^. To determine whether sweet GRNs are involved in taste-evoked warm preference, we silenced all sweet GRNs. We expressed the inwardly rectifying K^+^ channel Kir2.1 (*uas-Kir*) ^54^ using *Gr64f-Gal4*, which is expressed in all sweet GRNs in the proboscis and forelegs ^47,52,53^. Inactivation of all sweet GRNs showed a normal starvation response (Fig. 2C, white and gray bars, statistics shown as green stars). However, flies silencing all sweet GRNs failed to show a warm preference after 10 min of sucralose refeeding (Fig. 2C, blue bar, statistics shown as green and red stars, Table S1). This phenotype was similar to the data obtained with the sweet Gr mutant strains (Fig. 2A and 2B). On the other hand, control flies (*Gr64f-Gal4/+* and *uas-Kir/+*) showed a normal starvation response and a taste-evoked warm preference (Fig. 2D and 2E, gray and blue bars, respectively, statistics shown as green and red stars, Table S1). Thus, our data indicate that sweet GRNs are required for taste-evoked warm preference.

To further investigate whether activation of sweet GRNs induces a warm preference, we used the optogenetic approach, a red light sensitive channelrhodopsin, CsChrimson ^55,56^. Starved flies were given water containing 0.8 mM all-trans-retinal (ATR), the chromophore required for CsChrimson activation. These flies were not fed sucralose; instead, gustatory neurons in starved flies were excited by red light pulses (flashing on and off at 10 Hz) for 10 min (Fig. 2F). In this case, although the flies were not refed, the gustatory neurons were artificially excited by CsChrimson activation so that we could evaluate the effect of excitation of sweet GRNs on taste-evoked warm preference.

CsChrimson was expressed in sweet GRNs in the proboscis and legs (all sweet GRNs) using *Gr64f-Gal4*. These flies showed a normal starvation response (Fig. 2G-2J, white and gray bars, statistics shown as green and red stars, Table S1). Excitation of all sweet GRNs by red light pulses elicited a warm preference, and Tp was intermediate between fed and starved flies, suggesting partial recovery (Fig. 2G, yellow bar, statistics shown as green and red stars, Fig. S2B Table S1). However, neither excitation of Gr5a- (Fig. 2H) nor Gr64a-expressing neurons (Fig. 2I) induced a warm preference (yellow bars, Table S1). While *Gr64a-Gal4* is expressed only in the legs, *Gr5a-Gal4* is expressed in the proboscis and legs, but does not cover all sweet GRNs like *Gr64f-Gal4* ^52,53^. Notably, control flies (*UAS-CsChrimson/+*) did not show a warm preference to red light pulses with ATR application (Fig. 2J). Taken together, our data suggest that excitation of all sweet GRNs results in a warm preference.

We next asked whether the sweet Grs contribute to the nutrient-induced warm preference. We found that all these starved flies did not increase Tp after 10 min of glucose intake (Fig. 2A-2C, green bars, statistics shown as green and red stars, Table S1). All control flies showed normal responses to 10 min of glucose refeeding (Fig. 2D and 2E, green bars, Table S1). The data suggest that the sweet Grs which we tested are potentially expressed in tissues/neurons required for internal nutrient sensing. Notably, we found that flies increased Tp after 10 min of refeeding with fly food containing carbohydrate, fat, and protein (Fig. 2A-2C, orange bars, statistics shown as green and red stars, Table S1). All control flies showed normal responses to 10 min of fly food intake (Fig. 2D and 2E, orange bars, Table S1). The data suggest that gustatory neurons are required for warm preference in carbohydrate refeeding, but not for other nutrients such as fat or protein (see Discussion). Because flies have sensory neurons that detect fatty acids ^57–59^ or amino acids ^60–63^, these neurons may drive the response to fly food intake. This is likely why the sweet-insensitive flies can still recover after eating fly food (Fig 2A-2C, orange bars).

### The temperature-sensing neurons are involved in taste-evoked warm preference

The warm-sensing neurons, anterior cells (ACs), and the cold-sensing *R11F02-Gal4*-expressing neurons control temperature preference behavior ^26,44,64^. Small ectotherms such as *Drosophila* set their Tp to avoid noxious temperatures using temperature information from cold- and warm-sensing neurons ^17,18,44^. We have previously shown that starved flies choose a lower Tp, the so-called hunger-driven lower Tp ^26^. ACs control the hunger-driven lower Tp, but cold-sensing *R11F02-Gal4-*expressing neurons do not ^26^. ACs express transient receptor potential A1 (TrpA1), which responds to a warm temperature >25 °C ^44,65^. The set point of ACs in fed flies, which is ∼25 °C, is lowered in starved flies. Therefore, the lower set point of ACs corresponds to the lower Tp in starved flies.

First, we asked whether ACs are involved in taste-evoked warm preference. Because the ACs are important for the hunger-driven lower Tp ^26^, the AC-silenced flies did not show a significant difference in Tp between fed and starved conditions for only one overnight of starvation^26^. Therefore, we first extended the starvation time to two overnights so that the AC-silenced flies showed a significant difference in Tp between fed and starved conditions (Fig. 3A, white and gray bars, statistics shown as green and red stars, Table S1). Importantly, longer periods of starvation do not affect the ability of *w^1118^* flies to recover (Fig. S3).

**Figure 3.**
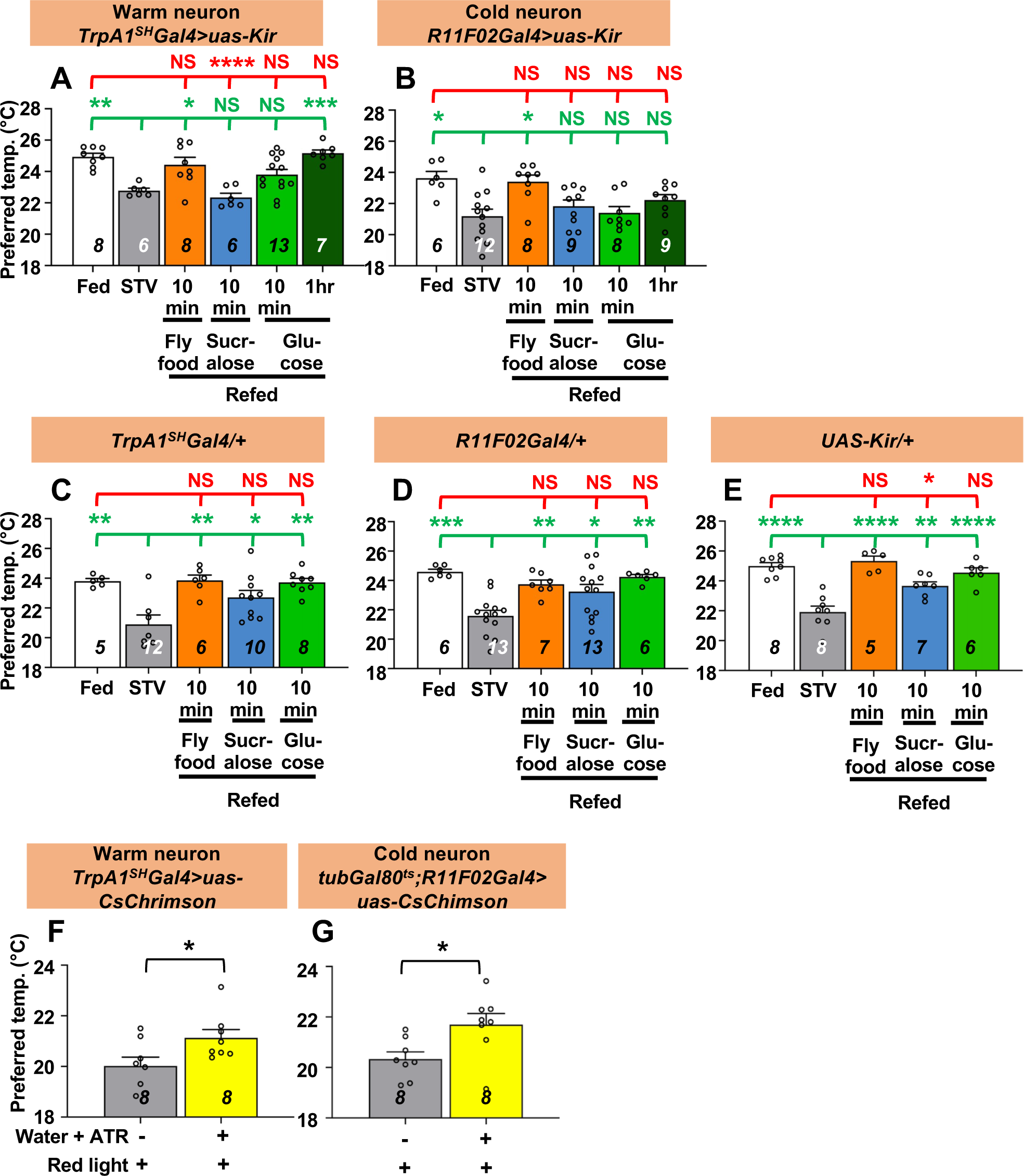
Both warm and cold temperature sensing neurons are involved in taste-evoked warm preference. **(A-E)** Comparison of preferred temperature (Tp) of flies between fed (white bar), starved (STV; gray bar), and refed (orange, blue, green, or dark green bar) conditions. Starvation was applied for 2 overnights (ON). Starved flies were refed with fly food for 10 min (orange bar), sucralose for 10 min (blue bar), or glucose for 10 min (green bar) or 1 hr (dark green bar). The Shapiro-Wilk test was used to test for normality. One-way ANOVA or Kruskal-Wallis test was used for statistical analysis. Red or green stars indicate Tukey’s post hoc test or Dunn’s test compared between each experiment and the fed (red) or starved (green) condition, respectively. **(F and G)** Comparison of Tp between starved (STV; gray bar) and all-trans-retinal (ATR; yellow bar) starved flies. Starvation was performed for 2ON. Warm neurons (F) or cold neurons (G) in starved flies expressed CsChrimson, which was excited by red light pulses for 10 min. The Shapiro-Wilk test was performed to test for normality. Student’s t-test or Kolmogorov-Smirnov test was used for statistical analysis. (G) *tubGal80^ts^; R11F02-Gal4>uas-CsChrimson* flies were reared at 18°C, and emerged adults were collected and stored at 29°C. See Materials and Methods for details. These behavioral experiments were performed on ZT4-7. The dots on each bar indicate individual Tp in the assays. Numbers in italics indicate the number of experiments. All data shown are means with SEM. *p < 0.05. **p < 0.01. ***p < 0.001. ****p < 0.0001. NS

To examine whether ACs regulate taste-evoked warm preference, we refed sucralose to AC-silenced flies for 10 min. We found that the Tp of the refed flies was still similar to that of the starved flies (Fig. 3A, blue bar, statistics shown as green and red stars, Table S1), indicating that sucralose refeeding could not restore Tp and that ACs are involved in taste-evoked warm preference. Significantly, even when the AC-silenced flies were starved for two overnights, they were able to recover Tp to the normal food for 10 min and to glucose for 1 hr (Fig. 3A, orange and green bars), suggesting that the starved AC-silenced flies were still capable of recovery.

Other temperature-sensing neurons involved in temperature preference behavior are cold-sensing *R11F02-Gal4-*expressing neurons ^26,64^. To determine whether *R11F02-Gal4-*expressing neurons are involved in taste-evoked warm preference, we silenced *R11F02-Gal4*-expressing neurons using *uas-Kir*. We found that the flies showed a significant difference in Tp between fed and starved conditions, but the flies did not show a warm preference upon sucralose refeeding (Fig. 3B, gray and blue bars, statistics shown as green and red stars, Table S1). As controls, *TrpA1^SH^-Gal4/*+, *R11F02-Gal4/*+ and *uas-Kir/*+ flies showed normal starvation response and taste-evoked warm preference (Fig. 3C-3E).

To further ensure the results, we used optogenetics to artificially excite warm and cold neurons with *TrpA1^SH^-Gal4* and *R11F02-Gal4*, respectively, by red light pulses for 10 min. We compared the Tp of starved flies with and without ATR under red light. Tp of starved flies with ATR was significantly increased (Fig. 3F and 3G, yellow bars, statistics shown as green and red stars, Table S1) compared to those without ATR (Fig. 3F and 3G, gray bars, Table S1). Therefore, these data indicate that ACs and *R11F02-Gal4-*expressing neurons are required for taste-evoked warm preference.

Next, we asked whether temperature-sensitive neurons contribute to the nutrient-induced warm preference. We used the warm- or cold neuron silenced flies (*TrpA1^SH^-Gal4* or *R11F02-Gal4>uas-Kir*) and found that all these starved flies did not increase Tp after 10 min glucose intake (Fig. 3A and 3B, green bars, Table S1), but increased Tp after 10 min refeeding with fly food containing carbohydrate, fat, and protein (Fig. 3A and 3B, orange bars, Table S1). Notably, AC-silenced flies increased Tp after 1 hr glucose intake (Fig. 3A, green bars, Table S1). All control flies showed normal responses to both 10 min glucose refeeding and fly food intake (Fig. 3C-3E, green bars, Table S1). The data suggest that temperature-sensing neurons are required for warm preference in carbohydrate refeeding, but not in other foods such as fat or protein (see Discussion).

### Olfaction is possibly involved in a warm preference for hungry flies

We also investigated the potential effects of olfaction. We used mutants of the odorant receptor co-receptor, *Orco* (*Orco^1^*), which has an olfactory defect ^66^. We found that the flies showed a significant difference in Tp between fed and starved conditions, but the flies did not show a warm preference upon sucralose refeeding (Fig. S4A and S4C, gray and blue bars, statistics shown as green and red stars, Table S1). We found that all of these starved flies increased Tp after 10 min of glucose or fly food intake and showed a full recovery after 1 hr of glucose intake (Fig. S4B and S4C, orange and green bars, statistics shown as green and red stars, Table S1). The data suggest that olfaction may be involved in the warm preference in sucralose refeeding.

### Internal state influences taste-evoked warm preference in hungry flies

Internal state strongly influences motivation to feed. However, how internal state influences starving animals to exhibit a food response remains unclear. Hunger represents the food-deficient state in the body, which induces the release of hunger signals such as neuropeptide Y (NPY). NPY promotes foraging and feeding behavior in mammals and flies ^67^. While intracerebroventricular injection of NPY induces the cephalic phase response (CPR) ^68^, injection of NPY antagonists suppresses CPR in dogs, suggesting that NPY is a regulator of CPR in mammals ^69^. Therefore, we first focused on neuropeptide F (NPF) and small neuropeptide F (sNPF), which are the *Drosophila* homologue and orthologue of mammalian NPY, respectively ^67^, and asked whether they are involved in taste-evoked warm preference. In *NPF* mutant (*NPF^−/−^*) or *sNPF* hypomorph (*sNPF hypo*) mutant, we found that the Tp of fed and starved flies were significantly different, showing a normal starved response (Fig. 4A and 4B: white and gray bars, statistics shown as green and red stars, Table S1). However, they failed to show a taste-evoked warm preference after 10 min of sucralose refeeding (Fig. 4A and 4B: blue bars, statistics shown as green and red stars, Table S1). Thus, NPF and sNPF are required for taste-evoked warm preference after sucralose refeeding.

**Figure 4.**
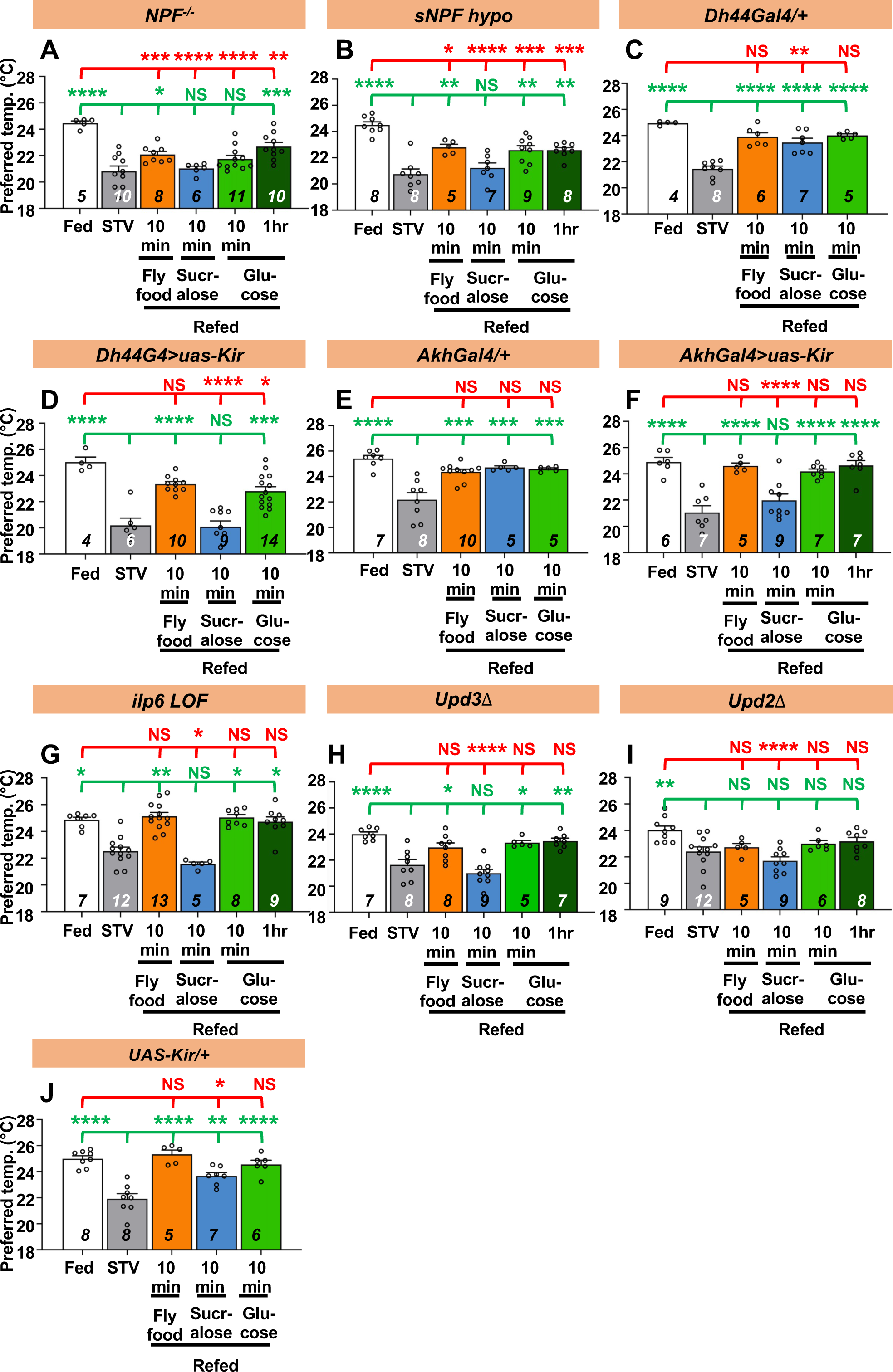
Hunger signals are involved in taste-evoked warm preference. Comparison of preferred temperature (Tp) of flies between fed (white bar), starved (STV; gray bar), and refed states (orange, blue, green, or dark green). Flies were starved for 2 overnights (ON) except for *ilp6* mutant flies (3ON). Starved flies were refed with fly food (orange bar), sucralose (blue bar), or glucose (green bar) for 10 min. These behavioral experiments were conducted on ZT4-7. Dots on each bar indicate individual Tp in the assays. Numbers in italics indicate the number of trials. Shapiro-Wilk test was performed for normality test. One-way ANOVA was used for statistical analysis. Red or green stars indicate Tukey’s post hoc test comparing between each experiment to the fed (red) or starved (green) condition, respectively. All data presented are means with SEM. *p < 0.05. **p < 0.01. ***p < 0.001. ****p < 0.0001. NS indicates not significant.

We next asked whether NPF and sNPF are involved in the nutrient-induced warm preference during glucose refeeding. We found that starved *NPF^−/−^* mutants significantly increased Tp after 1 hr of glucose refeeding (Fig. 4A, dark green bar, statistics shown as green and red stars, Table S1). We also found that starved *sNPF hypo* mutants significantly increased Tp after 10 min and 1 hr of glucose refeeding (Fig. 4B, green and dark green bars, statistics shown as green and red stars, Table S1). In addition, both *NPF^−/−^* and *sNPF hypo* mutants increased Tp after 10 min of fly food refeeding (Fig. 4A and 4B, orange bars, statistics shown as green and red stars, Table S1). However, when these flies were fed both glucose and fly food, they did not reach the same Tp as the fed flies. These data suggest that NPF and sNPF also play a role in the modulation of nutrient-induced warm preference.

### Factors involving hunger regulate taste-evoked warm preference

Based on the above results, we hypothesized that hunger signals might be involved in taste-evoked warm preference. To test this hypothesis, we focused on several factors involved in the hunger state. The major hunger signals ^24^, diuretic hormone 44 (DH44), and adipokinetic hormone (AKH) are the mammalian corticotropin-releasing hormone homologue ^70^ and the functional glucagon homologue, respectively ^71,72^. We found that DH44- or AKH-expressing neuron silenced flies failed to show a taste-evoked warm preference after sucralose refeeding. However, they were able to increase Tp after 10 min glucose or fly food refeeding (Fig. 4D and 4F, green and orange bars, statistics shown as green and red stars, Table S1), suggesting a normal nutrient-induced warm preference. The control flies (*Dh44-Gal4/+*, *Akh-Gal4/+*, and *UAS-Kir/+*) showed a normal warm preference to sucralose (Fig. 4C, 4E and 4J, blue bars, statistics shown as green and red stars, Table S1), glucose (Fig. 4C, 4E and 4J, green bars, statistics shown as green and red stars, Table S1), and normal fly food refeeding (Fig. 4C, 4E and 4J, orange bars, statistics shown as green and red stars, Table S1). The data suggest that DH44 or AKH neurons are required for taste-evoked warm preference, but not for nutrient-induced warm preference.

Insulin-like peptide 6 (Ilp6) is a homologue of mammalian insulin-like growth factor 1 (IGF1), and *ilp6* mRNA expression is increased in starved flies ^73–75^. Because Ilp6 is important for the hunger-driven lower Tp ^26^, the *ilp6* mutants did not show a significant difference in Tp between fed and starved conditions for only one overnight starvation. Therefore, we first extended the starvation time to three overnights. We found that the *ilp6* loss-of-function (*ilp6 LOF*) mutant failed to show a taste-evoked warm preference after sucralose refeeding (Fig. 4G, blue bars, statistics shown as green and red stars, Table S1), but did show a nutrient-induced warm preference after glucose refeeding (Fig. 4G, green bars, statistics shown as green and red stars, Table S1). These data suggest that Ilp6 is required for taste-evoked warm preference but not for nutrient-induced warm preference after glucose or fly food refeeding.

Unpaired3 (Upd3) is a *Drosophila* cytokine that is upregulated under nutritional stress ^76^. We found that the *upd3* mutants failed to show a taste-evoked warm preference after sucralose refeeding (Fig. 4H, blue bar, statistics shown as green and red stars, Table S1), but showed a nutrient-induced warm preference after glucose or normal food refeeding (Fig. 4H, green and orange bars, statistics shown as green and red stars, Table S1). These data suggest that Upd3 is required for taste-evoked warm preference, but not for nutrient-induced warm preference after glucose or normal food refeeding.

We also examined the role of the satiety factor Unpaired2 (Upd2), a functional leptin homologue in flies ^77^, in taste-evoked warm preference. The *upd2* mutants failed to show a warm preference after refeeding of sucralose, glucose or fly food (Fig. 4I, blue, green and orange bars, statistics shown as green and red stars, Table S1). Thus, our data suggest that factors involved in the hunger state are required for taste-evoked warm preference.

### Flies show a taste-evoked warm preference at all times of the day

Animals anticipate feeding schedules at a time of day that is tightly controlled by the circadian clock ^41^. Flies show a rhythmic feeding pattern: one peak in the morning ^50^ or two peaks in the morning and evening ^78^. Because food cues induce a warm preference, we wondered whether feeding rhythm and taste-evoked warm preference are coordinated. If so, they should show a parallel phenotype.

Since flies exhibit one of the circadian outputs, the temperature preference rhythm (TPR) ^21^, Tp gradually increases during the day and peaks in the evening. First, we tested starvation responses at Zeitgeber time (ZT)1-3, 4-6, 7-9, and 10-12 under light and dark (LD) conditions, with flies being offered only water for 24 h prior to the experiments at each time point. We found that both *w^1118^* and *yellow^1^ white^1^* (*y^1^w^1^*) flies had higher Tp in the fed state and lower Tp in the starved state at all times of the day (Fig. 5A and 5B: black and gray lines; Table S1). Next, we refed sucralose to starved flies at all time points tested and examined taste-evoked warm preference. While starved *y^1^w^1^* flies showed a taste-evoked warm preference at all time points (Fig. 5B1, gray and blue lines, Table S1), starved *w^1118^* flies showed a significant taste-evoked warm preference at ZT4-6 and 10-12 (Fig. 5A1, gray and blue lines, Table S1).

**Figure 5.**
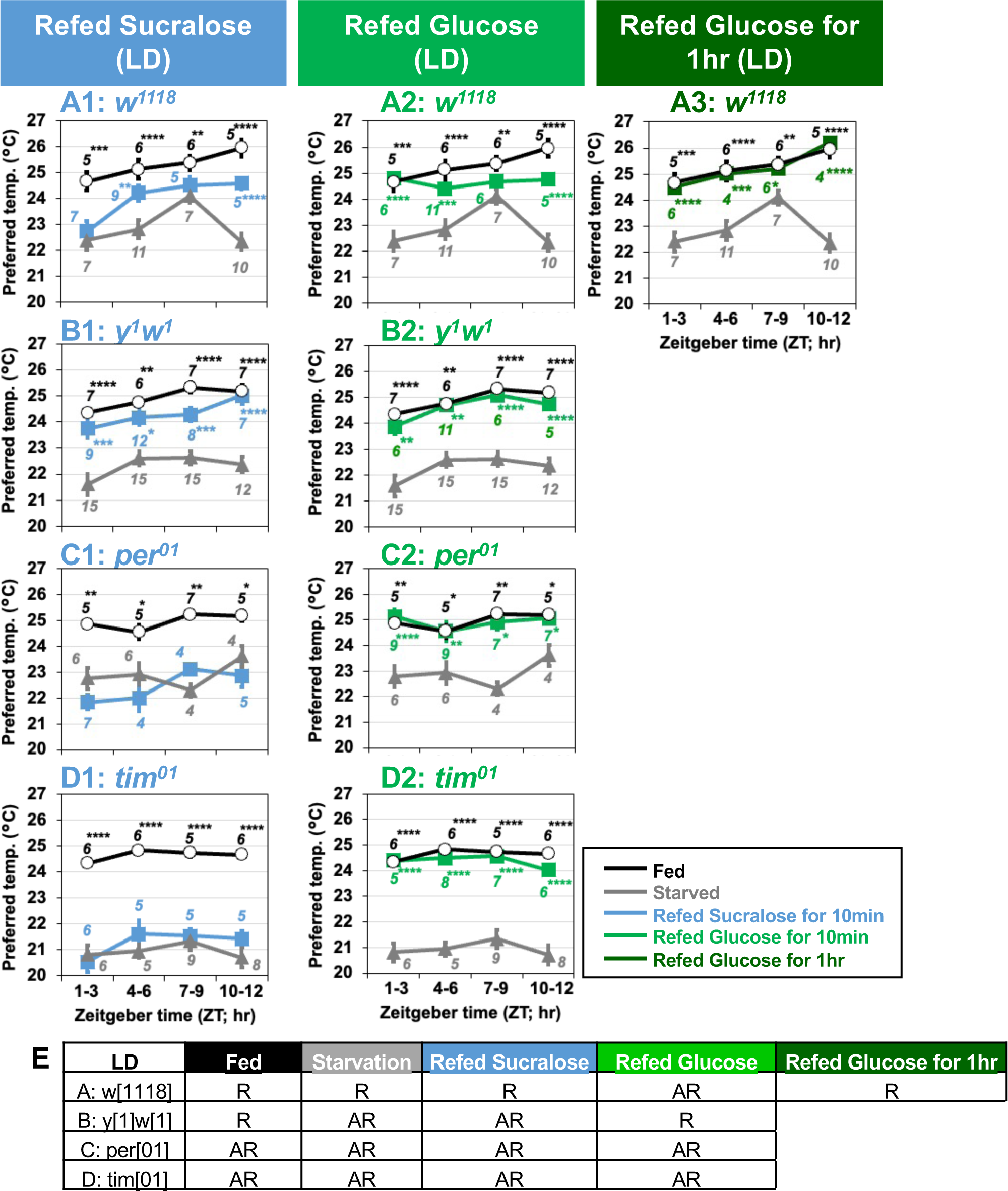
Clock genes are involved in taste-evoked warm preference. **(A-D)** Comparison of preferred temperature (Tp) of flies between fed (white circles), starved (STV; gray triangles), and refed (blue, green, or dark green squares) states. Flies were starved for 24 hrs. Starved flies were refed with sucralose (blue squares, A1-D1) or glucose for 10 min (green squares, A2-D2) or 1 hr (dark green squares, A3). After the 10 min or 1 hr refeeding, the temperature preference behavior assays were performed immediately at ZT1-3, ZT4-6, ZT7-9, and ZT10-12 in LD. The Shapiro-Wilk test was performed to test for normality. One-way ANOVA or Kruskal-Wallis test was used for statistical analysis. Stars indicate Tukey’s post hoc test or Dunn’s test compared between each experiment and the starved condition at the same time point. All data presented are means with SEM. *p<0.05. **p<0.01. ***p<0.001. ****p < 0.0001. **(E)** Comparison of Tp during the daytime in each feeding state. One-way ANOVA or Kruskal-Wallis test was used for statistical analysis between ZT1-3 and ZT7-9 or ZT10-12, respectively. R and AR indicate rhythmic and arrhythmic, respectively, during the daytime.

Because starved *w^1118^* flies showed an advanced phase shift of TPR with a peak at ZT7-9 (Fig. 5A, gray), it is likely that the highest Tp simply masks the taste-evoked warm preference at ZT7-9. We also focused on nutrient- (carbohydrate-) induced warm preference. Starved *w^1118^* and *y^1^w^1^* flies successfully increased Tp after 10 min of glucose refeeding (Fig. 5A2 and B2, green line, Table S1). Glucose refeeding for 1 hr resulted in Tp similar to that of fed *w^1118^* flies (Fig. 5A3, dark green line). Because the feeding rhythm peaks in the morning or morning/evening ^50,78^, our data suggest that the feeding rhythm and taste-evoked warm preference do not occur in parallel.

### Circadian clock genes are required for taste-evoked warm preference, but not for nutrient-induced warm preference

We asked whether the circadian clock is involved in taste-evoked warm preference. We used clock gene null mutants, *period^01^* (*per^01^*) and *timeless^01^* (*tim^01^*). Although they showed significant starvation responses (Fig. 5C and 5D, black and gray lines, Table S1), neither starved *per^01^* nor *tim^01^* mutants could show taste-evoked warm preference upon sucralose refeeding (Fig. 5C1 and 5D1, blue lines, Table S1). Nevertheless, they fully recovered upon glucose refeeding in LD at any time of day (Fig. 5C2 and 5D2, green lines, Table S1). Therefore, our data suggest that clock genes are required for taste-evoked warm preference, but not for nutrient-induced warm preference.

However, starved *per^01^* and *tim^01^* mutants may eat sucralose less frequently than glucose, which could result in a failure to show a taste-evoked warm preference. Therefore, we examined how often starved *per^01^* and *tim^01^* mutants touched glucose, sucralose, or water during the 30 min using FLIC assays ^50^ (Fig. S5A-S5C). Interestingly, starved *per^01^* and *tim^01^* mutants touched water significantly more often than glucose or sucralose (Fig. S5B and S5C, Table S1). Although starved *tim^01^* flies touched glucose slightly more than sucralose for only 10 min, this phenotype is not consistent with *per^01^* and *w^1118^* flies. However, these mutants still showed a similar Tp pattern for sucralose and glucose refeeding (Fig. 5C and 5D). The results suggest that although the *tim^01^* flies can eat sufficient amount of sucralose over glucose, their food intake does not affect the Tp behavioral phenotype. Thus, we conclude that in *per^01^* and *w^1118^* flies, the differential response between taste-evoked and nutrient-induced warm preferences is not due to feeding rate.

## DISCUSSION

When animals are hungry, sensory detection of food (sight, smell, or chewing) initiates digestion even before the food enters the stomach. The food-evoked responses are also observed in thermogenesis, heart rate, and respiratory rate in mammals ^14,15,79^. These responses are referred to as the cephalic phase response (CPR) and contribute to the physiological regulation of digestion, nutrient homeostasis, and daily energy homeostasis ^11^. While starved flies show a cold preference, we show here that the food cue, such as the excitation of gustatory neurons, triggers a warm preference, and the nutritional value triggers an even higher warm preference. Thus, when flies exit the starvation state, they use a two-step approach to recovery, taste-evoked and nutrient-induced warm preferences. The taste-evoked warm preference in *Drosophila* may be a physiological response potentially equivalent to CPR in mammals. Furthermore, we found that internal needs, controlled by hunger signals and circadian clock genes, influence taste-evoked warm preference. Thus, we propose that the taste-evoked response plays an important role in recovery and represents another layer of regulation of energy homeostasis.

### Tp is determined by the taste cue

Starved flies increase Tp in response to a nutrient-free taste cue (Fig. 1C and 1F), resulting in a taste-evoked warm preference. We showed that silencing of ACs or cold neurons caused a loss of taste-evoked warm preference (Fig. 3A-3E), and that excitation of ACs or cold neurons induced a taste-evoked warm preference (Fig. 3F and 3G). The data suggest that both warm and cold neurons are important for taste-evoked warm preference: while ACs are required for the hunger-driven lower Tp ^26^, both ACs and cold neurons are likely to be important for this taste-evoked warm preference.

The hunger-driven lower Tp is a slower response because starvation gradually lowers their Tp ^26^. In contrast, the taste-evoked warm preference is a rapid response. Once ACs and cold neurons are directly or indirectly activated, starved flies quickly move to a warmer area. This is interesting because even when ACs and cold neurons are activated by warm and cold, respectively, the activation of these neurons causes a warm preference. Given that the sensory detection of food (sight, smell, or chewing food) triggers CPR, the activation of these sensory neurons may induce CPR. Tp in sucralose-refed hungry flies is between that of fed and starved flies (Figs. 2 and 3), making it difficult to detect the smaller temperature differences using the calcium imaging experiments. Therefore, we speculate that both ACs and cold neurons may facilitate rapid recovery from starvation so that flies can quickly return to their preferred temperature - body temperature - to a normal state.

### Internal state influences taste-evoked warm preference

We show that mutants of genes involved in hunger and the circadian clock fail to show taste-evoked warm preference, suggesting that hunger and clock genes are important for taste-evoked warm preference. At a certain time of day, animals are hungry for food ^50,78^. Thus, the hungry state acts as a gatekeeper, opening the gate of the circuits when hungry flies detect the food information that leads to taste-evoked warm preference (Figs. 4 and S6, blue arrow). While most of the hunger signals we focused on are important for taste-evoked warm preference, some hunger signals are also required for both taste-evoked and nutrient-induced warm preferences (Fig. 4). Notably, sensory signals contribute to both taste-evoked and nutrient-induced warm preferences (Figs. 2, 3 and S6, blue and green arrows). Thus, taste-evoked warm preference and nutrient-induced warm preference differ at the internal state level, but not at the sensory level. This idea is analogous to appetitive memory formation. Sweet taste and nutrients regulate the different layers of the memory formation process. Recent evidence suggests that the rewarding process can be subdivided; the sweet taste is for short-term memory and the nutrient is for long-term memory ^80–82^. The data suggest that the taste-evoked response functions differently from the nutrient-induced response. Therefore, taste sensation is not just the precursor to nutrient sensing/absorption, but plays an essential role in the rapid initiation of a taste-evoked behavior that would help the animal survive.

### How do hunger signals or clock genes contribute to taste-evoked warm preference?

The hunger signaling hormones/peptides studied in this project are important for taste modulation. For example, mammalian NPY and its *Drosophila* homolog NPF modulate the output of taste signals ^49,83,84^. The AKH receptor is expressed in a subset of gustatory neurons that may modulate taste information for carbohydrate metabolism ^85^. Therefore, the hunger signals are likely to modulate the downstream of the sensory neurons, which may result in a taste-evoked warm preference.

Circadian clock genes control and coordinate the expression of many clock-controlled genes in the body ^86–89^. Therefore, we expect that the absence of clock genes will disrupt the molecular and neural networks of homeostasis, including metabolism, that are essential for animal life. For example, taste neurons express clock genes, and impaired clock function in taste neurons disrupts daily rhythms in feeding behavior ^90^. Temperature-sensing neurons transmit hot or cold temperature information to central clock neurons ^91–95^. Therefore, the disrupted central clock in clock mutants may respond imprecisely to temperature signals. There are many possible reasons why the lack of clock gene expression in the brain is likely to cause abnormal taste-evoked warm preference.

In addition, hunger signals may contribute to the regulation of circadian output. DH44 is located in the dorsomedial region of the fly brain, the pars intercerebralis (PI), and DH44-expressing neurons play a role in the output pathway of the central clock ^96,97^. Insulin-producing cells (IPCs) are also located in addition to DH44-expressing neurons ^98–101^. IPCs receive a variety of information, including circadian ^97,102^ and metabolic signals ^73–75,77^. and then transduce the signals downstream to release Ilps. Both Upd2 and Ilp6, which are expressed in the fat body respond to metabolic states and remotely regulate Ilp expression ^73–75,77^. Insect fat body is analogous to the fat tissues and liver in the vertebrates ^103,104^. Therefore, each hunger signal may have its specific function for taste-evoked warm preference. Further studies are needed to describe the entire process.

### Taste-evoked warm preference may be CPR in flies

The introduction of food into the body disrupts the internal milieu, so CPR is a necessary process that helps animals prepare for digestion. Specifically, in mammals, taste leads to an immediate increase in body temperature and metabolic rate. Starvation results in lower body temperatures, and chewing food, even before it enters the stomach, triggers a rapid increase in heat production, demonstrating CPR in thermogenesis ^14,15^.

Starved flies have a lower Tp ^26^. Because *Drosophila* is a small ectotherm, the lower Tp indicates a lower body temperature ^19,20^. Even when the flies do not receive food, the sweet taste and the excitation of sweet neurons induce starved flies to show a warm preference, which eventually leads to a warmer body temperature. In fact, CPR is known to be influenced by smell as well in mammals ^9^. We have shown in flies that olfactory mutants fail to show a warm preference when refed sucralose (Fig. S4). Starvation leads to lower body temperatures, and food cues, including taste and odor, rapidly induce a rise in body temperature before food enters the body. Thus, the taste-evoked warm preference in *Drosophila* may be a physiological response equivalent to one of the CPRs observed in mammals.

### CPR is essential because both starved mammals and starved flies must rapidly regulate their body temperature to survive

As soon as starved flies taste food, the sensory signals trigger CPR. They can move to a warmer place to prepare to raise their body temperature (Fig. S6, blue arrows). CPR may allow flies to choose a more hospitable place to restore their physiological state and allow for a higher metabolism, and eventually move on to the next step, such as foraging and actively seeking a mate before competitors arrive. Thus, CPR may be a strategy for the fly’s survival. Similarly, starvation or malnutrition in mammals leads to lower body temperatures ^6,7,105^, and biting food triggers heat production, which is CPR ^14–16^. Thus, while starvation in both flies and mammals leads to lower body temperatures, food cues initiate CPR by increasing body temperature and nutrient intake, resulting in full recovery from starvation. Our data suggest that *Drosophila* CPR may be a physiological response equivalent to CPR observed in other animals. Thus, *Drosophila* may shed new light on the regulation of CPR and provide a deeper understanding of the relationship between CPR and metabolism.

## Acknowledgements

We are grateful to Drs. Anupama Dahanukar, Hubert Amrein, Greg Suh, and Paul A. Garrity and the Bloomington Drosophila Fly Stock Center for the fly lines. We thank members of the Hamada laboratory for critical comments and advice on the manuscript, Dr. Richard A. Lang and members of his laboratory for comments and kindly sharing reagents, Dr. Satoshi Namekawa for kindly sharing reagents, and Matthew Batie and Nathan T. Petts for design and construction of the behavioral assays and red-light illumination apparatus. This research was supported by a RIP funding from Cincinnati Children’s Hospital, JST (Japan Science and Technology)/Precursory Research for Embryonic Science and Technology (PRESTO), the March of Dimes, and NIH R01 grant GM107582, NIH R21 grant NS112890, NIH R35 GM152154 grant, and NIH R34 grant NS132843 to F.N.H. S.H.S. is a Latin American Fellow in the Biomedical Sciences supported by the Pew Charitable Trusts and by NIH K99 grant NS133470. Research in the laboratory of J.C.C. is supported by NIH R01 grant DK124068.

## Contributions

F.N.H and Y.U designed the experiments. Y.U, E.N., T.N. and J.S. performed the temperature preference behavioral assays and data analysis. Y.U., S.H.S., S.S.U. and J.C. performed the feeding experiments and the data analysis. F.N.H and Y.U wrote the manuscript.

---

**Figure S1.**
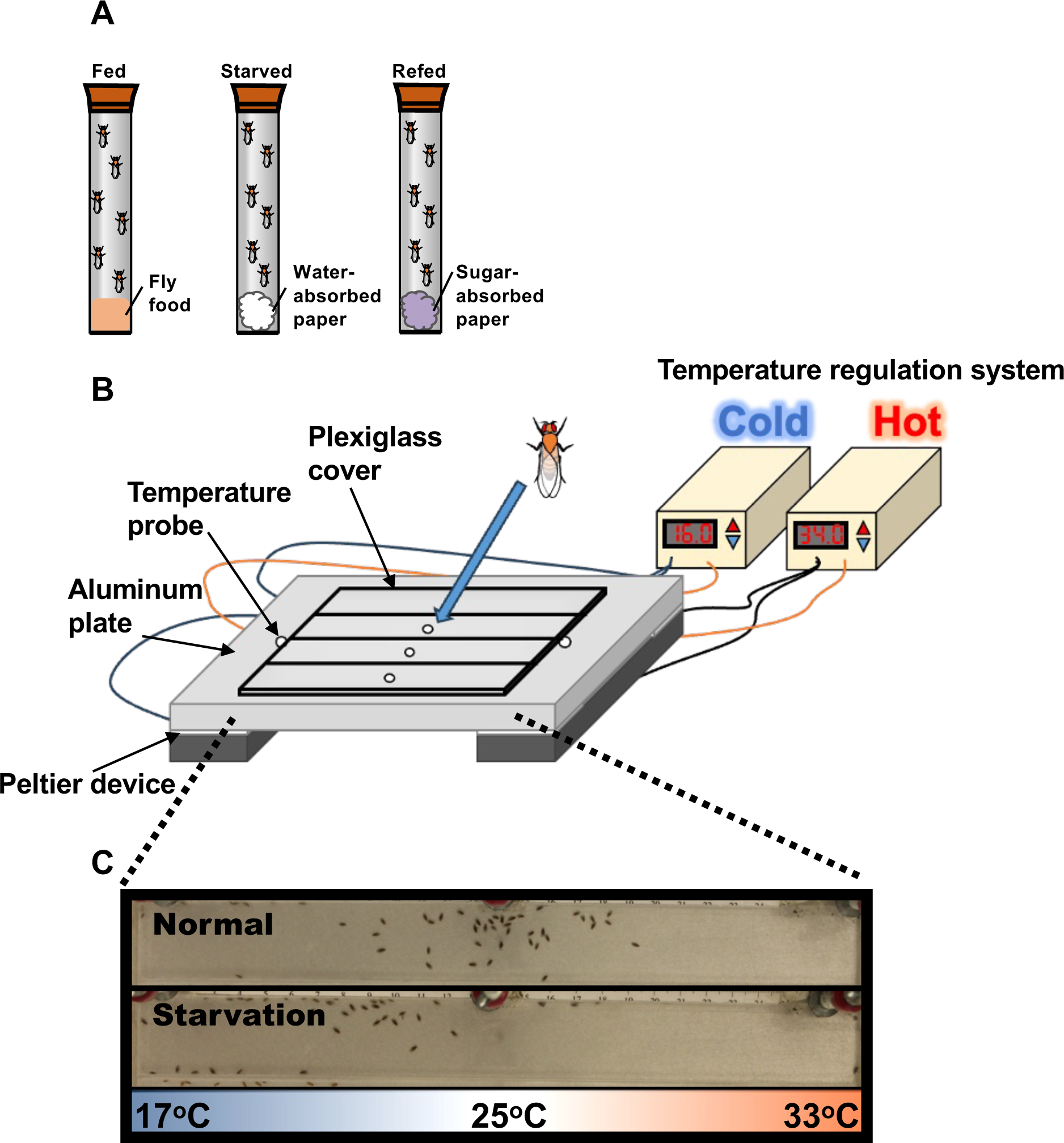
*Drosophila* temperature preference behavior assay (Related to Fig. 1) (A) Three different feeding conditions, fed, starved and refed, were used in this study. (B) A temperature gradient of 17-33°C in air is established in the chamber between the aluminum metal plate and the Plexiglas cover. Flies are applied through the hole into the chamber. Once the flies are applied, they spread out in the chamber and then gradually accumulate at the specific temperature ranges within 30 min, which is called the preferred temperature (Tp). (C) One of the representative results of Tp experiments (top, normal (fed flies); bottom, starvation (starved flies)). Since their body temperature is close to the temperature of their surrounding microenvironment, their body temperature is determined by measuring their Tp.

**Figure S2.**
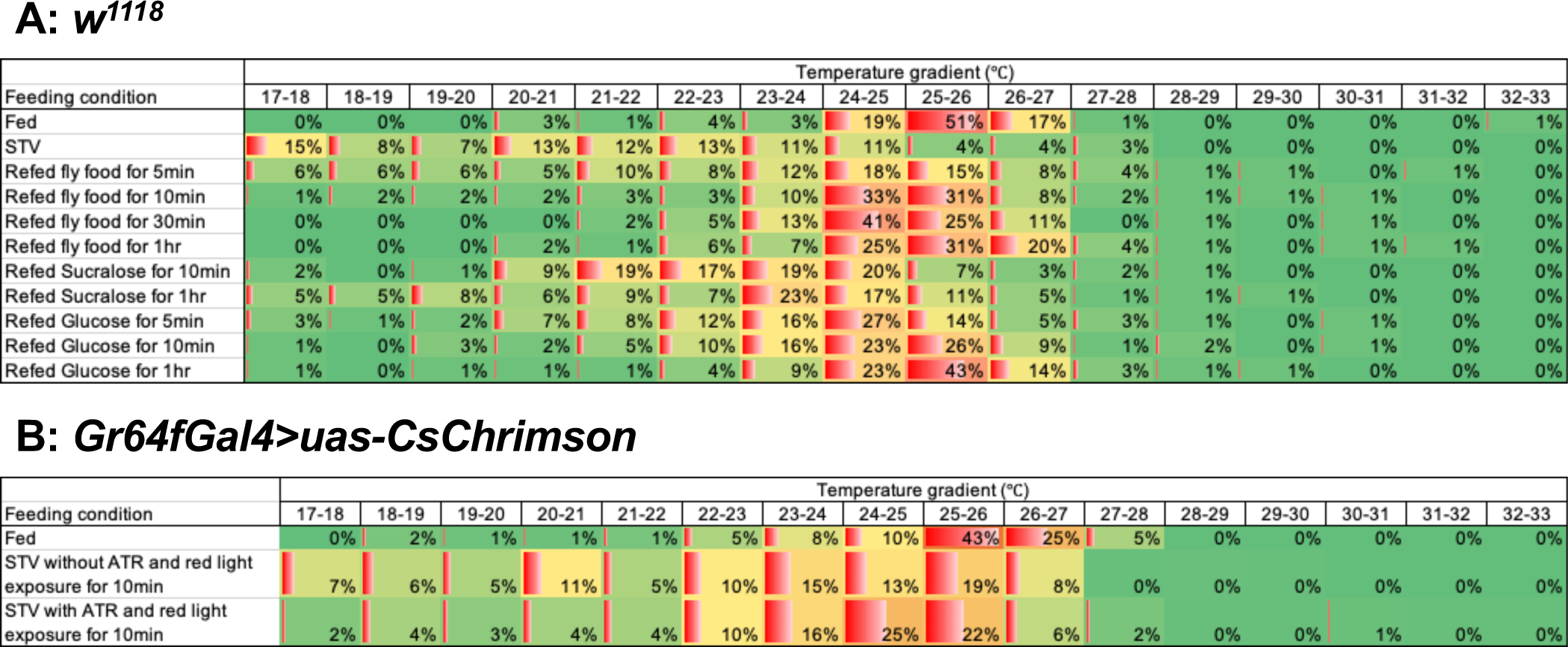
The distribution of temperature preference in each experimental condition (Related to Fig. 1 and 2) Comparison of the distribution of temperature preference in each feeding condition in *w^1118^* control flies (A) and the *Gr64fGal4>CsChrimson* flies (B). Heat maps were generated using the Conditional formatting tool in Microsoft Excel. The averaged percentages of flies settling on the apparatus within each one-degree temperature interval were used to create the heat maps. Each parameter to draw them using Conditional formatting tool is as follows: Minimum Value: 0, Midpoint Value: 15%, and Maximum Value: 60% for *w^1118^*. Minimum Value: 0, Midpoint Value: 10%, and Maximum Value: 45% for *Gr64fGal4>CsChrimson*.

**Figure S3.**
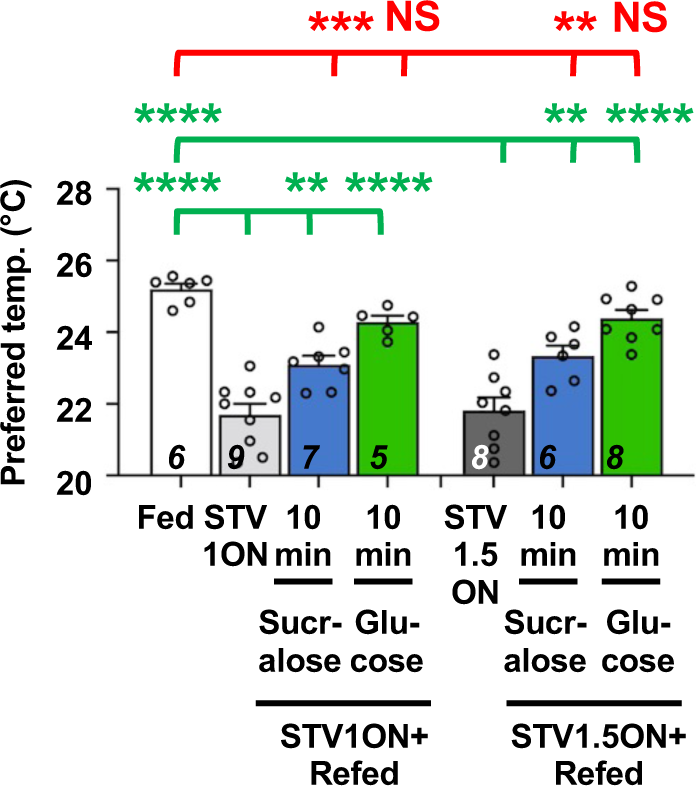
Duration of starvation is unlikely to affect the ability to recover (Related to Fig. 3) Comparison of preferred temperature (Tp) of *w^1118^* control flies between fed (white bar), starved (STV; gray bar), and refed conditions (blue or green bar). Flies were starved for overnight (STV1ON; gray bar) or 1.5 ON (STV1.5ON; dark gray bar), i.e. starved for 18-21 hrs or 26-29 hrs, respectively. Starved flies were refed with sucralose for 10 min (blue bar) or glucose for 10 min (green bar). These behavioral experiments were performed on ZT4-7. Dots on each bar indicate individual Tp in the assays. Numbers in italics indicate the number of trials. Shapiro-Wilk test was performed to test for normality. One-way ANOVA or Kruskal-Wallis test was used for statistical analysis. Red or green stars indicate Tukey’s post hoc test or Dunn’s test comparing differences between experimental and fed or starved conditions, respectively. All data shown are means with sem. *p < 0.05. **p < 0.01. ***p < 0.001. ****p < 0.0001. NS indicates no significance. The same data from Fed, STV1ON, STV1ON+refed sucralose for 10 min and STV1ON+refed sucralose for 10 min in Fig. 1B-F are used in Fig. S3.

**Figure S4.**
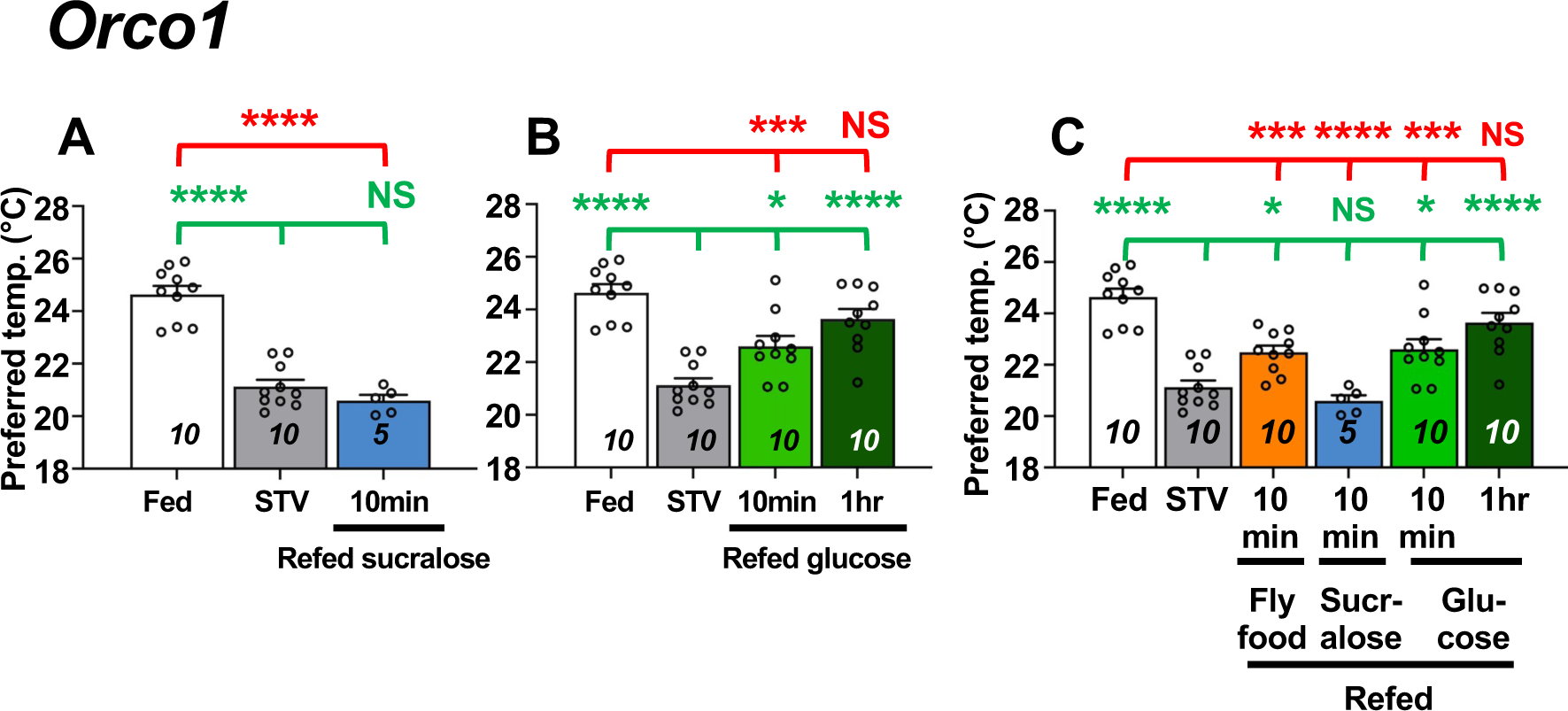
*Orco* is involved in the taste-evoked warm preference. Comparison of the preferred temperature (Tp) of *Orco*^1^ olfaction-deficient mutant flies between fed (white bar), starved (STV; gray bar), and refed (blue or green bar). *Orco*^1^ flies were starved for 2 overnights (ON). Starved flies were refed with sucralose for 10 min (blue bar; A) or glucose for 10 min (green bar; B). Pooled data of both sucralose- and glucose-refeeding bioassay data are shown in C. The Shapiro-Wilk test was performed to test for normality. One-way ANOVA or Kruskal-Wallis test was used for statistical analysis. Red or green stars indicate Tukey’s post hoc test or Dunn’s test compared between each experiment and the fed (red) or starved (green) condition, respectively.

**Figure S5.**
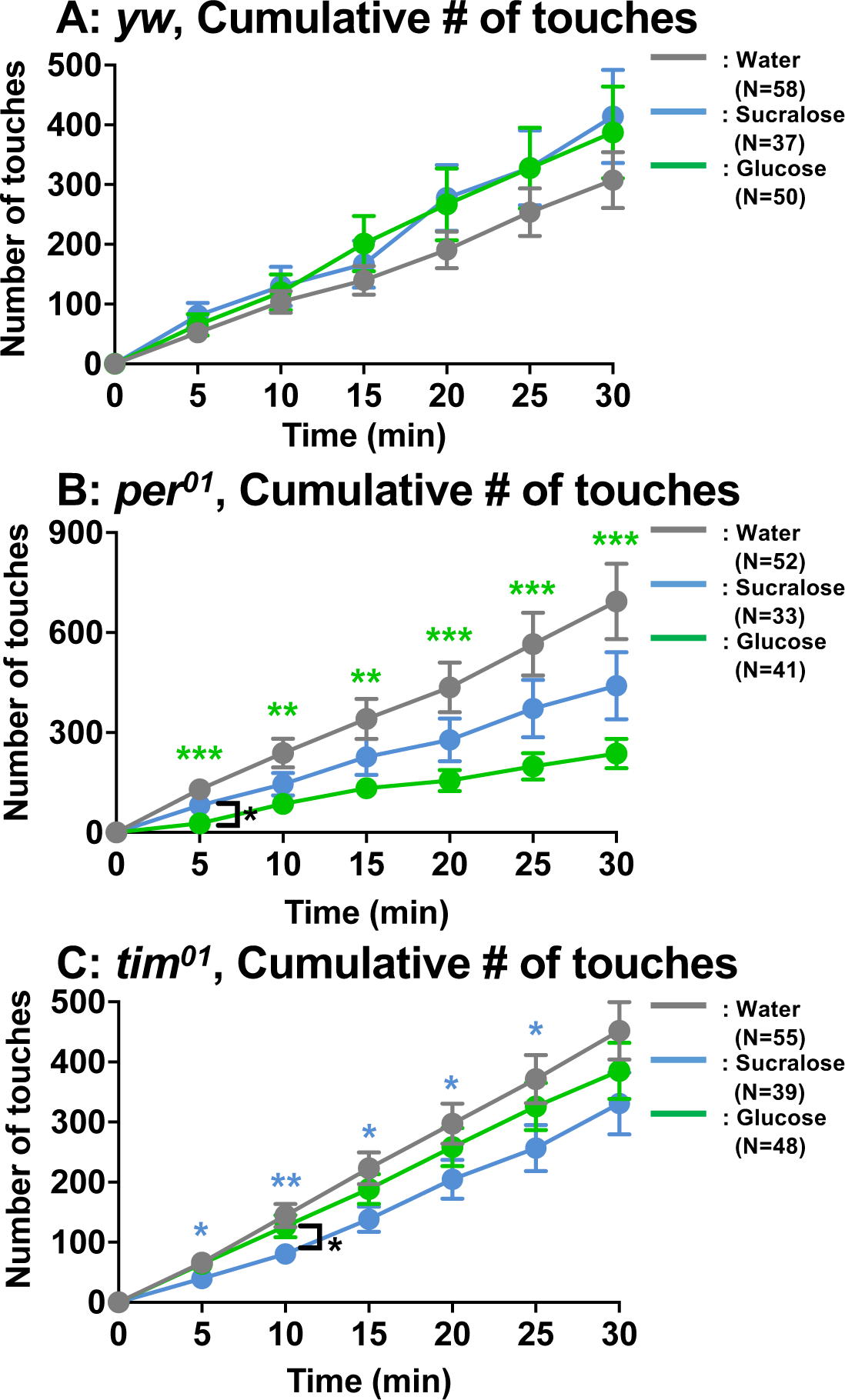
Number of touches to water, sucralose and glucose using *yw*, *per^01^* and *tim^01^* flies. (Related to Figures 1 and 5) Feeding assay: The number of touches to water (gray), 2.8 mM (equivalent to 5%) glucose (green), or 2.8 mM sucralose solution (blue) was examined using *yw*, *per^01^*, or *tim^01^* flies starved for 24 hrs. Cumulative number of touches for 0-30 min was plotted. Two-way ANOVA was used for multiple comparisons. Blue and green stars show Fisher’s LSD post hoc test comparing sucralose (blue stars) or glucose (green stars) solution feeding to water drinking. Black stars show the statistical difference between sucralose and glucose refeeding. All data presented are means with SEM. *p < 0.05. *F*p < 0.01. ***p < 0.001. ****p < 0.0001.

**Figure S6.**
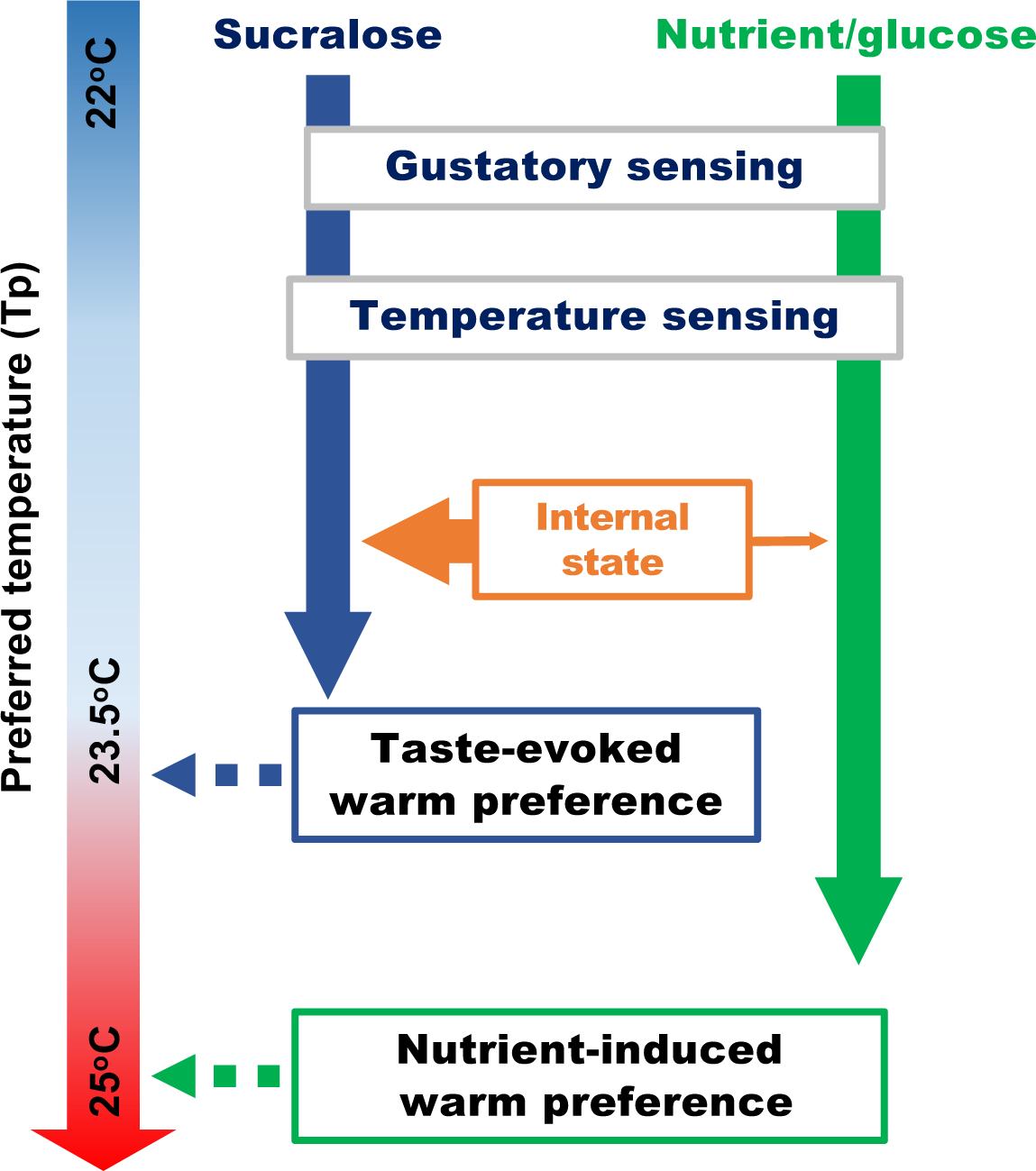
MODEL. A schematic diagram of the recovery of preferred temperature (Tp) by taste-evoked warm preference (blue arrows) and nutrient-induced warm preference (green arrows).

## Materials and Methods

All flies were reared under 12 hr-light/12 hr-dark (LD) cycles at 25°C and 60-70% humidity in an incubator (DRoS33SD, Powers Scientific Inc.) with an electric timer (light on: 8am; light off: 8pm). The light intensity was 1000-1400 lux. All flies were reared on custom fly food recipe, with the following composition per 1 L of food: 6.0 g sucrose, 7.3 g agar, 44.6 g cornmeal, 22.3 g yeast and 16.3 mL molasses, as previously described ^26^. *white^1118^* (*w^1118^*) and *yellow*^1^ *white*^1^ (*y^1^w*^1^) flies were used as control flies. *EP5Δ; Gr64a*^1^ (*Gr5a^−/−^; Gr64a^−/−^*) was kindly provided by Dr. Anupama Dahanukar ^47^. *R1; Gr5a-LexA; +; ΔGr61a, ΔGr64a-f* (*Gr5a^−/−^; Gr61a^−/−^, Gr64a-f^−/−^*) were kindly provided by Dr. Hubert Amrein ^51,52^. *Dh44-Gal4* was kindly provided by Dr. Greg Suh ^48^. *TrpA1^SH^-gal4* was kindly provided by Dr. Paul A. Garrity ^44^. Other fly lines were provided by Bloomington Drosophila stock center and Vienna Drosophila Stock Center.

### Temperature preference behavioral assay

The temperature preference behavior assays were examined using a temperatures gradient, set from 16-34°C and were performed for 30 min in an environmental room maintained at 25°C /60%-70% humidity, as previously described ^26^. Because starved flies showed a lower preferred temperature (Tp), the temperature regulation was lower than usual (Umezaki et al. 2018 Current Biology). We prepared a total of 40-50 flies (male and female flies were mixed) for fed condition experiments and 90-100 flies for overnight(s) starved and refed condition experiments for one trial. Flies were never reused in subsequent trials. In the starved and refed experiments, we prepared twice the number of flies needed for a trial because almost half of them died from starvation stress. Others climbed on the wall and ceiling (starved flies are usually hyperactive ^106^, even though we applied slippery coating chemicals (PTFE; Cat# 665800, Sigma or byFormica PTFE Plus, https://byformica.com/products/fluon-plus-ptfe-escape-prevention-coating) to the Plexiglass covers.

Behavioral assays were performed for 30 min at Zeitgeber time (ZT) 4-7 (light on and light off are defined as ZT0 and ZT12, respectively), and starvation was initiated at ZT9-10 (starved for 1, 2 or 3 O/N are 18-21 hr, 42-45 hr or 66-69 hr, respectively). As for STV1.5 for Fig. S3, starvation was initiated at ZT1-2 (26-29 hr). For the starvation assays, the collected flies were maintained on our fly food for at least one day and then transferred to plastic vials containing 3 mL of distilled water, which was absorbed by a Kimwipe paper. For refeeding experiments, starved flies were transferred to plastic vials containing 2 mL of 2.8 mM sugar solution (sucralose water/glucose water). Sugar solution is absorbed by half the size of a Kimwipe. The details of the starvation period are described in the following section (see “Starvation Condition”).

After the 30-min behavioral assay, the number of flies whose bodies were completely located on the aluminum plate was counted. Flies whose bodies were partially or completely located on the walls of the Plexiglass cover were not included in the data analysis. The percentage of flies within each one-degree temperature interval on the apparatus was calculated by dividing the number of flies within each one-degree interval by the total number of flies on the apparatus. The location of each one-degree interval was determined by measuring the temperature at 6 different points on the bottom of the apparatus. Data points were plotted as the percentage of flies within a one-degree temperature interval. The weighted mean of Tp was calculated by summing the product of the percentage of flies within a one-degree temperature interval and the corresponding temperature (e.g., fractional number of flies x 17.5°C + fractional number of flies x 18.5°C +……… fractional number of flies x 32.5°C). If the SEM of the averaged Tp was not < 0.3 after the five trials, additional trials were performed approximately 10 times until the SEM was < 0.3.

Microsoft Excel (Home tab>Conditional formatting tool>3-color scales and data bars) was used to create heat maps to show the distribution of flies in each experimental condition. The averaged percentages of flies that settled on the apparatus within each one-degree temperature interval were used to create the heat maps. Each scale value is as follows; minimum value: 0, midpoint value: 15%, and maximum value: 60% for *w^1118^*. Minimum value: 0, middle value: 10%, and maximum value: 45% for Gr64fGal4>CsChrimson.

### Starvation conditions and recovery

Most of the flies were starved for 2 overnights (O/N). Because some flies (e.g. *ilp6* mutants) show starvation resistance and seem to be still healthy even after 2 O/N of starvation. We had to starve them for 3 O/N to show a significant difference in Tp between fed and starved flies. On the other hand, some flies (e.g., *w^1118^* flies) are very sick after 3 O/N of starvation, in which case we only had to starve them for 1 day. Therefore, the starvation conditions we used for this manuscript are from 1 to 3 O/N.

First, we confirmed the starvation period by focusing on Tp which resulted in a statistically significant Tp difference between fed and starved flies; as mentioned above, flies prefer lower temperatures when starvation is prolonged ^26^. Therefore, when Tp was not statistically different between fed and starved flies, we extended the starvation period from 1 to 3 O/N. Importantly, we show in Fig. S3 that the duration of starvation does not affect the recovery effect. Furthermore, *w^1118^* flies cannot survive 42-49 or 66-69 hours of starvation.

### Temperature preference rhythm (TPR) assay

For the TPR assays, we performed temperature preference behavior assays in different time windows during the daytime (zeitgeber time (ZT) or circadian time (CT) 1-3, 4-6, 7-9, and 10-12) as described previously ^21,107^. Because starvation duration directly affects flies’ Tp ^26^ starvation was initiated at each time window to adjust the starvation duration at each time point, which means flies were starved for 24 hrs or 48 hrs but not 1 or 2 O/N. Each behavioral assay was not examined during these time periods (ZT or CT0-1 and 11.5-12) because of large phenotype variation around light on and light off.

Furthermore, insulin levels were shown to peak at 10 min and gradually decline ^108^. Also, how quickly the flies can consume food is unclear. These factors may influence temperature preference behavior. Therefore, to minimize these effects, we decided to test the temperature preference behavioral assay immediately after the flies had eaten the food.

### Optogenetic activation

For the optogenetic activation of the target neurons for behavioral assays, the red-light-sensitive channelrodopsin, *UAS-CsChrimson*, was crossed with each Gal4 driver. Flies were reared on fly food at 25°C and 60-70% humidity under LD cycles in an incubator (DRoS33SD, Powers Scientific Inc.) with an electric timer. After the flies emerged, adult flies were collected and maintained on fly food for 1-2 days. The next day, flies were transferred to water with or without 0.8 mM all-trans retinal (ATR; #R2500, Sigma) diluted in dimethyl sulfoxide (DMSO; #472301, Sigma) for 2 O/N. To activate flies expressing UAS-CsChrimson crossed with Gal4 drivers, we used a 627-nm red light-emitting diode (LED) equipped with a pulsed photoillumination system (10 Hz, 0.08 mWmm-2). Flies were exposed to pulsed red light for 10 min, which corresponds to the refeeding period. This photoillumination system was used in an incubator (Sanyo Scientific, MIR-154) and followed by temperature preference behavioral assays.

The *R11F02-Gal4>uas-CsChrimson* flies do not develop into adults and die in the pupal stage. Therefore, the Gal4/Gal80ts system was used to restrict *uas-CsChrimson* expression. The *Gal80^ts^* is a temperature sensitive allele of *Gal80* that causes *Gal4* inhibition at 18°C and activation at 29°C ^109^.The *tubGal80^ts^; R11F02-Gal4>uas-CsChrimson* flies were reared on fly food at 18°C. Emerging adult flies were collected and kept on fly food at 29°C. The next day, flies were transferred to water (starved condition) with or without 0.8 mM ATR for 2 O/N. Starved flies with or without ATR application were exposed to pulsed red light for 10 min (equivalent to the refeeding period) and then immediately loaded into the behavioral apparatus for behavioral assays to measure their Tp. All flies were treated with ATR after they had fully developed into the adults. This means that Gal4-expressing cells were activated by red light via CsChrimson only at adult stages.

### Feeding assay

To measure individual fly feeding, we used the Fly Liquid Interaction Counter (FLIC) system ^50^. Groups of 1-2 day old male and female flies were starved for 24 hours starting between ZT4-5 (12-1pm). Individual flies were then loaded into FLIC monitors. Flies were acclimated to the monitors for 30 min with access to water in the feeding wells. At the start of the feeding study, water was replaced with 2.8 mM sucralose or 2.8 mM glucose solution (equivalent to 5% glucose concentration) and the number of licks (touches) was recorded for 30 min. Water, sucralose, or glucose water was administered individually in separate experiments. Assays were performed on ZT4-7. To account for the potential confounding effect of startle response when food is changed, wells where water was replaced with new water were used as a control. FLIC raw data were analyzed using the FLIC R code master (Pletcher Lab, available at https://github.com/PletcherLab/FLIC_R_Code). Lick counts were obtained and summed in 5-min window bins, while cumulative licks were obtained by successively summing the licks in these bins.

## REFERENCES

1. Root, C.M., Ko, K.I., Jafari, A., and Wang, J.W. (2011). Presynaptic facilitation by neuropeptide signaling mediates odor-driven food search. Cell 145, 133–144. 10.1016/j.cell.2011.02.008.

2. Kain, P., and Dahanukar, A. (2015). Secondary taste neurons that convey sweet taste and starvation in the Drosophila brain. Neuron 85, 819–832. 10.1016/j.neuron.2015.01.005.

3. Soria-Gomez, E., Bellocchio, L., Reguero, L., Lepousez, G., Martin, C., Bendahmane, M., Ruehle, S., Remmers, F., Desprez, T., Matias, I., et al. (2014). The endocannabinoid system controls food intake via olfactory processes. Nat Neurosci 17, 407–415. 10.1038/nn.3647.

4. Hanci, D., and Altun, H. (2016). Hunger state affects both olfactory abilities and gustatory sensitivity. Eur Arch Otorhinolaryngol 273, 1637–1641. 10.1007/s00405-015-3589-6.

5. Keys, A., Brožek, J., Henschel, A., Mickelsen, O., and Taylor, H.L. (1950). The Biology of Human Starvation. University of Minnesota Press.

6. Piccione, G., Caola, G., and Refinetti, R. (2002). Circadian modulation of starvation-induced hypothermia in sheep and goats. Chronobiology international 19, 531–541.

7. Sakurada, S., Shido, O., Sugimoto, N., Hiratsuka, Y., Yoda, T., and Kanosue, K. (2000). Autonomic and behavioural thermoregulation in starved rats. The Journal of physiology 526 Pt 2, 417–424.

8. Smeets, P.A., Erkner, A., and de Graaf, C. (2010). Cephalic phase responses and appetite. Nutr Rev 68, 643–655. 10.1111/j.1753-4887.2010.00334.x.

9. Chen, Y., and Knight, Z.A. (2016). Making sense of the sensory regulation of hunger neurons. Bioessays 38, 316–324. 10.1002/bies.201500167.

10. Power, M.L., and Schulkin, J. (2008). Anticipatory physiological regulation in feeding biology: cephalic phase responses. Appetite 50, 194–206. 10.1016/j.appet.2007.10.006.

11. Zafra, M.A., Molina, F., and Puerto, A. (2006). The neural/cephalic phase reflexes in the physiology of nutrition. Neuroscience and biobehavioral reviews 30, 1032–1044. 10.1016/j.neubiorev.2006.03.005.

12. Pavlov, I.P. (1902). The Work of the Digestive Glands. London : Charles Griffin.

13. Garrison, J.L., and Knight, Z.A. (2017). Linking smell to metabolism and aging. Science 358, 718–719. 10.1126/science.aao5474.

14. LeBlanc, J. (2000). Nutritional implications of cephalic phase thermogenic responses. Appetite 34, 214–216. 10.1006/appe.1999.0283.

15. LeBlanc, J., and Cabanac, M. (1989). Cephalic postprandial thermogenesis in human subjects. Physiol Behav 46, 479–482.

16. LeBlanc, J., Cabanac, M., and Samson, P. (1984). Reduced postprandial heat production with gavage as compared with meal feeding in human subjects. The American journal of physiology 246, E95–101. 10.1152/ajpendo.1984.246.1.E95.

17. Sayeed, O., and Benzer, S. (1996). Behavioral genetics of thermosensation and hygrosensation in Drosophila. Proceedings of the National Academy of Sciences of the United States of America 93, 6079–6084.

18. Dillon, M.E., Wang, G., Garrity, P.A., and Huey, R.B. (2009). Review: Thermal preference in Drosophila. J Therm Biol 34, 109–119. 10.1016/j.jtherbio.2008.11.007.

19. Stevenson, R.D. (1985). Body size and limits to the daily range of body temperature in terrestrial ectotherms. Am. Nat., 102–117.

20. Stevenson, R.D. (1985). The relative importance of behavioral and physiological adjustments controlling body temperature in terrestrial ectotherms. The American Naturalist 126.

21. Kaneko, H., Head, L.M., Ling, J., Tang, X., Liu, Y., Hardin, P.E., Emery, P., and Hamada, F.N. (2012). Circadian Rhythm of Temperature Preference and Its Neural Control in Drosophila. Current biology : CB 22, 1851–1857. S0960-9822(12)00934-7 [pii] 10.1016/j.cub.2012.08.006.

22. Goda, T., and Hamada, F.N. (2019). Drosophila Temperature Preference Rhythms: An Innovative Model to Understand Body Temperature Rhythms. Int J Mol Sci 20. 10.3390/ijms20081988.

23. Goda, T., Umezaki, Y., and Hamada, F.N. (2023). Molecular and Neural Mechanisms of Temperature Preference Rhythm in Drosophila melanogaster. Journal of biological rhythms 38, 326–340. 10.1177/07487304231171624.

24. Lin, S., Senapati, B., and Tsao, C.H. (2019). Neural basis of hunger-driven behaviour in Drosophila. Open Biol 9, 180259. 10.1098/rsob.180259.

25. Zhang, D.W., Xiao, Z.J., Zeng, B.P., Li, K., and Tang, Y.L. (2019). Insect Behavior and Physiological Adaptation Mechanisms Under Starvation Stress. Frontiers in physiology 10, 163. 10.3389/fphys.2019.00163.

26. Umezaki, Y., Hayley, S.E., Chu, M.L., Seo, H.W., Shah, P., and Hamada, F.N. (2018). Feeding-State-Dependent Modulation of Temperature Preference Requires Insulin Signaling in Drosophila Warm-Sensing Neurons. Current biology : CB 28, 779–787.e773. 10.1016/j.cub.2018.01.060.

27. Berrigan, D., and Partridge, L. (1997). Influence of temperature and activity on the metabolic rate of adult Drosophila melanogaster. Comp Biochem Physiol A Physiol 118, 1301–1307.

28. Schilman, P.E., Waters, J.S., Harrison, J.F., and Lighton, J.R. (2011). Effects of temperature on responses to anoxia and oxygen reperfusion in Drosophila melanogaster. The Journal of experimental biology 214, 1271–1275. 10.1242/jeb.052357.

29. Ja, W.W., Carvalho, G.B., Mak, E.M., de la Rosa, N.N., Fang, A.Y., Liong, J.C., Brummel, T., and Benzer, S. (2007). Prandiology of Drosophila and the CAFE assay. Proceedings of the National Academy of Sciences of the United States of America 104, 8253–8256. 10.1073/pnas.0702726104.

30. Itskov, P.M., Moreira, J.M., Vinnik, E., Lopes, G., Safarik, S., Dickinson, M.H., and Ribeiro, C. (2014). Automated monitoring and quantitative analysis of feeding behaviour in Drosophila. Nature communications 5, 4560. 10.1038/ncomms5560.

31. Wang, G.H., and Wang, L.M. (2019). Recent advances in the neural regulation of feeding behavior in adult Drosophila. J Zhejiang Univ Sci B 20, 541–549. 10.1631/jzus.B1900080.

32. Pool, A.H., and Scott, K. (2014). Feeding regulation in Drosophila. Curr Opin Neurobiol 29, 57–63. 10.1016/j.conb.2014.05.008.

33. Snell, N.J., Fisher, J.D., Hartmann, G.G., Zolyomi, B., Talay, M., and Barnea, G. (2022). Complex representation of taste quality by second-order gustatory neurons in Drosophila. Current biology : CB 32, 3758–3772 e3754. 10.1016/j.cub.2022.07.048.

34. Kim, H., Kirkhart, C., and Scott, K. (2017). Long-range projection neurons in the taste circuit of Drosophila. Elife 6. 10.7554/eLife.23386.

35. Liu, W.W., Mazor, O., and Wilson, R.I. (2015). Thermosensory processing in the Drosophila brain. Nature 519, 353–357. 10.1038/nature14170.

36. Frank, D.D., Jouandet, G.C., Kearney, P.J., Macpherson, L.J., and Gallio, M. (2015). Temperature representation in the Drosophila brain. Nature 519, 358–361. 10.1038/nature14284.

37. Cedernaes, J., Waldeck, N., and Bass, J. (2019). Neurogenetic basis for circadian regulation of metabolism by the hypothalamus. Genes Dev 33, 1136–1158. 10.1101/gad.328633.119.

38. Longo, V.D., and Panda, S. (2016). Fasting, Circadian Rhythms, and Time-Restricted Feeding in Healthy Lifespan. Cell Metab 23, 1048–1059. 10.1016/j.cmet.2016.06.001.

39. Hirayama, M., Mure, L., S, and Panda, S. (2018). Circadian regulation of energy intake in mammals. Current Opinion in Physiology 5, 141–148.

40. Greenhill, C. (2018). Benefits of time-restricted feeding. Nat Rev Endocrinol 14, 626. 10.1038/s41574-018-0093-2.

41. Chaix, A., Manoogian, E.N.C., Melkani, G.C., and Panda, S. (2019). Time-Restricted Eating to Prevent and Manage Chronic Metabolic Diseases. Annu Rev Nutr 39, 291–315. 10.1146/annurev-nutr-082018-124320.

42. Blasiak, A., Gundlach, A.L., Hess, G., and Lewandowski, M.H. (2017). Interactions of Circadian Rhythmicity, Stress and Orexigenic Neuropeptide Systems: Implications for Food Intake Control. Front Neurosci 11, 127. 10.3389/fnins.2017.00127.

43. Panda, S. (2016). Circadian physiology of metabolism. Science 354, 1008–1015. 10.1126/science.aah4967.

44. Hamada, F.N., Rosenzweig, M., Kang, K., Pulver, S.R., Ghezzi, A., Jegla, T.J., and Garrity, P.A. (2008). An internal thermal sensor controlling temperature preference in Drosophila. Nature 454, 217–220. nature07001 [pii] 10.1038/nature07001.

45. Biolchini, M., Murru, E., Anfora, G., Loy, F., Banni, S., Crnjar, R., and Sollai, G. (2017). Fat storage in Drosophila suzukii is influenced by different dietary sugars in relation to their palatability. PloS one 12, e0183173. 10.1371/journal.pone.0183173.

46. Park, J.H., Carvalho, G.B., Murphy, K.R., Ehrlich, M.R., and Ja, W.W. (2017). Sucralose Suppresses Food Intake. Cell Metab 25, 484–485. 10.1016/j.cmet.2017.02.011.

47. Dahanukar, A., Lei, Y.T., Kwon, J.Y., and Carlson, J.R. (2007). Two Gr genes underlie sugar reception in Drosophila. Neuron 56, 503–516. 10.1016/j.neuron.2007.10.024.

48. Dus, M., Min, S., Keene, A.C., Lee, G.Y., and Suh, G.S. (2011). Taste-independent detection of the caloric content of sugar in Drosophila. Proceedings of the National Academy of Sciences of the United States of America 108, 11644–11649. 10.1073/pnas.1017096108.

49. Wang, Q.P., Lin, Y.Q., Zhang, L., Wilson, Y.A., Oyston, L.J., Cotterell, J., Qi, Y., Khuong, T.M., Bakhshi, N., Planchenault, Y., et al. (2016). Sucralose Promotes Food Intake through NPY and a Neuronal Fasting Response. Cell Metab 24, 75–90. 10.1016/j.cmet.2016.06.010.

50. Ro, J., Harvanek, Z.M., and Pletcher, S.D. (2014). FLIC: high-throughput, continuous analysis of feeding behaviors in Drosophila. PloS one 9, e101107. 10.1371/journal.pone.0101107.

51. Yavuz, A., Jagge, C., Slone, J., and Amrein, H. (2014). A genetic tool kit for cellular and behavioral analyses of insect sugar receptors. Fly 8, 189–196. 10.1080/19336934.2015.1050569.

52. Fujii, S., Yavuz, A., Slone, J., Jagge, C., Song, X., and Amrein, H. (2015). Drosophila sugar receptors in sweet taste perception, olfaction, and internal nutrient sensing. Current biology : CB 25, 621–627. 10.1016/j.cub.2014.12.058.

53. Thoma, V., Knapek, S., Arai, S., Hartl, M., Kohsaka, H., Sirigrivatanawong, P., Abe, A., Hashimoto, K., and Tanimoto, H. (2016). Functional dissociation in sweet taste receptor neurons between and within taste organs of Drosophila. Nature communications 7, 10678. 10.1038/ncomms10678.

54. Baines, R.A., Uhler, J.P., Thompson, A., Sweeney, S.T., and Bate, M. (2001). Altered electrical properties in Drosophila neurons developing without synaptic transmission. The Journal of neuroscience : the official journal of the Society for Neuroscience 21, 1523–1531.

55. Klapoetke, N.C., Murata, Y., Kim, S.S., Pulver, S.R., Birdsey-Benson, A., Cho, Y.K., Morimoto, T.K., Chuong, A.S., Carpenter, E.J., Tian, Z., et al. (2014). Independent optical excitation of distinct neural populations. Nat Methods 11, 338–346. 10.1038/nmeth.2836.

56. Simpson, J.H., and Looger, L.L. (2018). Functional Imaging and Optogenetics in Drosophila. Genetics 208, 1291–1309. 10.1534/genetics.117.300228.

57. Masek, P., and Keene, A.C. (2013). Drosophila fatty acid taste signals through the PLC pathway in sugar-sensing neurons. PLoS genetics 9, e1003710. 10.1371/journal.pgen.1003710.

58. Ahn, J.E., Chen, Y., and Amrein, H. (2017). Molecular basis of fatty acid taste in Drosophila. Elife 6. 10.7554/eLife.30115.

59. Brown, E.B., Shah, K.D., Palermo, J., Dey, M., Dahanukar, A., and Keene, A.C. (2021). Ir56d-dependent fatty acid responses in Drosophila uncover taste discrimination between different classes of fatty acids. Elife 10. 10.7554/eLife.67878.

60. Croset, V., Schleyer, M., Arguello, J.R., Gerber, B., and Benton, R. (2016). A molecular and neuronal basis for amino acid sensing in the Drosophila larva. Sci Rep 6, 34871. 10.1038/srep34871.

61. Ganguly, A., Pang, L., Duong, V.K., Lee, A., Schoniger, H., Varady, E., and Dahanukar, A. (2017). A Molecular and Cellular Context-Dependent Role for Ir76b in Detection of Amino Acid Taste. Cell Rep 18, 737–750. 10.1016/j.celrep.2016.12.071.

62. Chen, Y.D., and Dahanukar, A. (2017). Molecular and Cellular Organization of Taste Neurons in Adult Drosophila Pharynx. Cell Rep 21, 2978–2991. 10.1016/j.celrep.2017.11.041.

63. Steck, K., Walker, S.J., Itskov, P.M., Baltazar, C., Moreira, J.M., and Ribeiro, C. (2018). Internal amino acid state modulates yeast taste neurons to support protein homeostasis in Drosophila. Elife 7. 10.7554/eLife.31625.

64. Ni, L., Klein, M., Svec, K.V., Budelli, G., Chang, E.C., Ferrer, A.J., Benton, R., Samuel, A.D., and Garrity, P.A. (2016). The Ionotropic Receptors IR21a and IR25a mediate cool sensing in Drosophila. Elife 5. 10.7554/eLife.13254.

65. Tang, X., Platt, M.D., Lagnese, C.M., Leslie, J.R., and Hamada, F.N. (2013). Temperature integration at the AC thermosensory neurons in Drosophila. The Journal of neuroscience : the official journal of the Society for Neuroscience 33, 894–901. 10.1523/JNEUROSCI.1894-12.2013.

66. Larsson, M.C., Domingos, A.I., Jones, W.D., Chiappe, M.E., Amrein, H., and Vosshall, L.B. (2004). Or83b encodes a broadly expressed odorant receptor essential for Drosophila olfaction. Neuron 43, 703–714. 10.1016/j.neuron.2004.08.019.

67. Nassel, D.R., and Wegener, C. (2011). A comparative review of short and long neuropeptide F signaling in invertebrates: Any similarities to vertebrate neuropeptide Y signaling? Peptides 32, 1335–1355. 10.1016/j.peptides.2011.03.013.

68. Geoghegan, J.G., Lawson, D.C., Cheng, C.A., Opara, E., Taylor, I.L., and Pappas, T.N. (1993). Intracerebroventricular neuropeptide Y increases gastric and pancreatic secretion in the dog. Gastroenterology 105, 1069–1077. 10.1016/0016-5085(93)90951-8.

69. Lee, M.C., Lawson, D.C., and Pappas, T.N. (1994). Neuropeptide Y functions as a physiologic regulator of cephalic phase acid secretion. Regul Pept 52, 227–234. 10.1016/0167-0115(94)90057-4.

70. Dus, M., Lai, J.S., Gunapala, K.M., Min, S., Tayler, T.D., Hergarden, A.C., Geraud, E., Joseph, C.M., and Suh, G.S. (2015). Nutrient Sensor in the Brain Directs the Action of the Brain-Gut Axis in Drosophila. Neuron 87, 139–151. 10.1016/j.neuron.2015.05.032.

71. Lee, G., and Park, J.H. (2004). Hemolymph sugar homeostasis and starvation-induced hyperactivity affected by genetic manipulations of the adipokinetic hormone-encoding gene in Drosophila melanogaster. Genetics 167, 311–323.

72. Kim, S.K., and Rulifson, E.J. (2004). Conserved mechanisms of glucose sensing and regulation by Drosophila corpora cardiaca cells. Nature 431, 316–320. 10.1038/nature02897.

73. Okamoto, N., Yamanaka, N., Yagi, Y., Nishida, Y., Kataoka, H., O’Connor, M.B., and Mizoguchi, A. (2009). A fat body-derived IGF-like peptide regulates postfeeding growth in Drosophila. Dev Cell 17, 885–891. 10.1016/j.devcel.2009.10.008.

74. Slaidina, M., Delanoue, R., Gronke, S., Partridge, L., and Leopold, P. (2009). A Drosophila insulin-like peptide promotes growth during nonfeeding states. Dev Cell 17, 874–884. 10.1016/j.devcel.2009.10.009.

75. Bai, H., Kang, P., and Tatar, M. (2012). Drosophila insulin-like peptide-6 (dilp6) expression from fat body extends lifespan and represses secretion of Drosophila insulin-like peptide-2 from the brain. Aging Cell 11, 978–985. 10.1111/acel.12000.

76. Woodcock, K.J., Kierdorf, K., Pouchelon, C.A., Vivancos, V., Dionne, M.S., and Geissmann, F. (2015). Macrophage-derived upd3 cytokine causes impaired glucose homeostasis and reduced lifespan in Drosophila fed a lipid-rich diet. Immunity 42, 133–144. 10.1016/j.immuni.2014.12.023.

77. Rajan, A., and Perrimon, N. (2012). Drosophila cytokine unpaired 2 regulates physiological homeostasis by remotely controlling insulin secretion. Cell 151, 123–137. 10.1016/j.cell.2012.08.019.

78. Xu, K., Zheng, X., and Sehgal, A. (2008). Regulation of feeding and metabolism by neuronal and peripheral clocks in Drosophila. Cell Metab 8, 289–300. 10.1016/j.cmet.2008.09.006.

79. Nederkoorn, C., Smulders, F.T., and Jansen, A. (2000). Cephalic phase responses, craving and food intake in normal subjects. Appetite 35, 45–55. 10.1006/appe.2000.0328.

80. Musso, P.Y., Lampin-Saint-Amaux, A., Tchenio, P., and Preat, T. (2017). Ingestion of artificial sweeteners leads to caloric frustration memory in Drosophila. Nature communications 8, 1803. 10.1038/s41467-017-01989-0.

81. Gruber, F., Knapek, S., Fujita, M., Matsuo, K., Bracker, L., Shinzato, N., Siwanowicz, I., Tanimura, T., and Tanimoto, H. (2013). Suppression of conditioned odor approach by feeding is independent of taste and nutritional value in Drosophila. Current biology : CB 23, 507–514. 10.1016/j.cub.2013.02.010.

82. Huetteroth, W., Perisse, E., Lin, S., Klappenbach, M., Burke, C., and Waddell, S. (2015). Sweet taste and nutrient value subdivide rewarding dopaminergic neurons in Drosophila. Current biology : CB 25, 751–758. 10.1016/j.cub.2015.01.036.

83. Inagaki, H.K., Jung, Y., Hoopfer, E.D., Wong, A.M., Mishra, N., Lin, J.Y., Tsien, R.Y., and Anderson, D.J. (2014). Optogenetic control of Drosophila using a red-shifted channelrhodopsin reveals experience-dependent influences on courtship. Nat Methods 11, 325–332. 10.1038/nmeth.2765.

84. Herness, S., and Zhao, F.L. (2009). The neuropeptides CCK and NPY and the changing view of cell-to-cell communication in the taste bud. Physiol Behav 97, 581–591. 10.1016/j.physbeh.2009.02.043.

85. Bharucha, K.N., Tarr, P., and Zipursky, S.L. (2008). A glucagon-like endocrine pathway in Drosophila modulates both lipid and carbohydrate homeostasis. The Journal of experimental biology 211, 3103–3110. 10.1242/jeb.016451.

86. Giebultowicz, J.M. (2018). Circadian regulation of metabolism and healthspan in Drosophila. Free Radic Biol Med 119, 62–68. 10.1016/j.freeradbiomed.2017.12.025.

87. Mazzoccoli, G., Pazienza, V., and Vinciguerra, M. (2012). Clock genes and clock-controlled genes in the regulation of metabolic rhythms. Chronobiology international 29, 227–251. 10.3109/07420528.2012.658127.

88. Lin, Y., Han, M., Shimada, B., Wang, L., Gibler, T.M., Amarakone, A., Awad, T.A., Stormo, G.D., Van Gelder, R.N., and Taghert, P.H. (2002). Influence of the period-dependent circadian clock on diurnal, circadian, and aperiodic gene expression in Drosophila melanogaster. Proceedings of the National Academy of Sciences of the United States of America 99, 9562–9567. 10.1073/pnas.132269699.

89. Bozek, K., Relogio, A., Kielbasa, S.M., Heine, M., Dame, C., Kramer, A., and Herzel, H. (2009). Regulation of clock-controlled genes in mammals. PloS one 4, e4882. 10.1371/journal.pone.0004882.

90. Chatterjee, A., Tanoue, S., Houl, J.H., and Hardin, P.E. (2010). Regulation of gustatory physiology and appetitive behavior by the Drosophila circadian clock. Current biology : CB 20, 300–309. 10.1016/j.cub.2009.12.055.

91. Tang, X., Roessingh, S., Hayley, S.E., Chu, M.L., Tanaka, N.K., Wolfgang, W., Song, S., Stanewsky, R., and Hamada, F.N. (2017). The role of PDF neurons in setting preferred temperature before dawn in Drosophila. Elife 6. 10.7554/eLife.23206.

92. Alpert, M.H., Frank, D.D., Kaspi, E., Flourakis, M., Zaharieva, E.E., Allada, R., Para, A., and Gallio, M. (2020). A Circuit Encoding Absolute Cold Temperature in Drosophila. Current biology : CB 30, 2275–2288.e2275. 10.1016/j.cub.2020.04.038.

93. Marin, E.C., Büld, L., Theiss, M., Sarkissian, T., Roberts, R.J.V., Turnbull, R., Tamimi, I.F.M., Pleijzier, M.W., Laursen, W.J., Drummond, N., et al. (2020). Connectomics Analysis Reveals First-, Second-, and Third-Order Thermosensory and Hygrosensory Neurons in the Adult Drosophila Brain. Current biology : CB 30, 3167–3182.e3164. 10.1016/j.cub.2020.06.028.

94. Yadlapalli, S., Jiang, C., Bahle, A., Reddy, P., Meyhofer, E., and Shafer, O.T. (2018). Circadian clock neurons constantly monitor environmental temperature to set sleep timing. Nature. 10.1038/nature25740.

95. Jin, X., Tian, Y., Zhang, Z.C., Gu, P., Liu, C., and Han, J. (2021). A subset of DN1p neurons integrates thermosensory inputs to promote wakefulness via CNMa signaling. Current biology : CB 31, 2075–2087 e2076. 10.1016/j.cub.2021.02.048.

96. King, A.N., Barber, A.F., Smith, A.E., Dreyer, A.P., Sitaraman, D., Nitabach, M.N., Cavanaugh, D.J., and Sehgal, A. (2017). A Peptidergic Circuit Links the Circadian Clock to Locomotor Activity. Current biology : CB 27, 1915–1927 e1915. 10.1016/j.cub.2017.05.089.

97. Barber, A.F., Fong, S.Y., Kolesnik, A., Fetchko, M., and Sehgal, A. (2021). Drosophila clock cells use multiple mechanisms to transmit time-of-day signals in the brain. Proceedings of the National Academy of Sciences of the United States of America 118. 10.1073/pnas.2019826118.

98. Cao, C., and Brown, M.R. (2001). Localization of an insulin-like peptide in brains of two flies. Cell and tissue research 304, 317–321. 10.1007/s004410100367.

99. Brogiolo, W., Stocker, H., Ikeya, T., Rintelen, F., Fernandez, R., and Hafen, E. (2001). An evolutionarily conserved function of the Drosophila insulin receptor and insulin-like peptides in growth control. Current biology : CB 11, 213–221.

100. Ikeya, T., Galic, M., Belawat, P., Nairz, K., and Hafen, E. (2002). Nutrient-dependent expression of insulin-like peptides from neuroendocrine cells in the CNS contributes to growth regulation in Drosophila. Current biology : CB 12, 1293–1300.

101. Rulifson, E.J., Kim, S.K., and Nusse, R. (2002). Ablation of insulin-producing neurons in flies: growth and diabetic phenotypes. Science 296, 1118–1120. 10.1126/science.1070058.

102. Cavanaugh, D.J., Geratowski, J.D., Wooltorton, J.R., Spaethling, J.M., Hector, C.E., Zheng, X., Johnson, E.C., Eberwine, J.H., and Sehgal, A. (2014). Identification of a circadian output circuit for rest:activity rhythms in Drosophila. Cell 157, 689–701. 10.1016/j.cell.2014.02.024.

103. Li, S., Yu, X., and Feng, Q. (2019). Fat Body Biology in the Last Decade. Annu Rev Entomol 64, 315–333. 10.1146/annurev-ento-011118-112007.

104. Arrese, E.L., and Soulages, J.L. (2010). Insect fat body: energy, metabolism, and regulation. Annu Rev Entomol 55, 207–225. 10.1146/annurev-ento-112408-085356.

105. Cintron-Colon, R., Sanchez-Alavez, M., Nguyen, W., Mori, S., Gonzalez-Rivera, R., Lien, T., Bartfai, T., Aid, S., Francois, J.C., Holzenberger, M., and Conti, B. (2017). Insulin-like growth factor 1 receptor regulates hypothermia during calorie restriction. Proceedings of the National Academy of Sciences of the United States of America 114, 9731–9736. 10.1073/pnas.1617876114.

106. Yang, Z., Yu, Y., Zhang, V., Tian, Y., Qi, W., and Wang, L. (2015). Octopamine mediates starvation-induced hyperactivity in adult Drosophila. Proceedings of the National Academy of Sciences of the United States of America 112, 5219–5224. 10.1073/pnas.1417838112.

107. Goda, T., Leslie, J.R., and Hamada, F.N. (2014). Design and analysis of temperature preference behavior and its circadian rhythm in Drosophila. Journal of visualized experiments : JoVE, e51097. 10.3791/51097.

108. Tsao, D.D., Chang, K.R., Kockel, L., Park, S., and Kim, S.K. (2023). A genetic strategy to measure insulin signaling regulation and physiology in Drosophila. PLoS genetics 19, e1010619. 10.1371/journal.pgen.1010619.

109. Southall, T.D., and Brand, A.H. (2008). Generation of Driver and Reporter Constructs for the GAL4 Expression System in Drosophila. CSH Protoc 2008, pdb prot5029. 10.1101/pdb.prot5029.

