## Supplemental Table for "Taste triggers a homeostatic temperature control in hungry flies"

### Table S1. Statistical analysis for preferred temperatures (Tp).

Fig. 1

Fig. 1B-F

| *$w^{1118}$* | | |
|---|---|---|
| Comparison of Tp between | | p value |
| Fed vs | Starvation | **** |
| | Refed fly food for 5 min | **** |
| | Refed fly food for 10 min | ns |
| | Refed fly food for 30 min | ns |
| | Refed fly food for 1 hr | ns |
| | Refed Sucralose for 10 min | *** |
| | Refed Sucralose for 1 hr | **** |
| | Refed Glucose for 10 min | ns |
| | Refed Glucose for 1 hr | ns |
| | Refed Fructose for 10 min | ns |
| | Refed Fructose for 1 hr | ns |
| Starvation vs | Refed fly food for 5 min | *** |
| | Refed fly food for 10 min | **** |
| | Refed fly food for 30 min | **** |
| | Refed fly food for 1 hr | **** |
| | Refed Sucralose for 10 min | * |
| | Refed Sucralose for 1 hr | * |
| | Refed Glucose for 10 min | **** |
| | Refed Glucose for 1 hr | **** |
| | Refed Fructose for 10 min | **** |
| | Refed Fructose for 1 hr | **** |

| | |
|---|---|
| p value | p<0.0001 |
| alpha | 0.05 |
| Multiple test (ANOVA and Tukey's post hoc test or Kruskal-Wallis test and Dunn's test) | Tukey test |
| F value (F (DFn, DFd)) | F (12, 75)=16.24 |

Fig. 1G

| *$w^{1118}$*, Cumulative # of licking | | |
|---|---|---|
| Time | Starvation vs Refed Sucralose | Starvation vs Refed Glucose |
| 0-5 | ** | * |

| | | |
|---|---|---|
| 5-10 | ** | * |
| 10-15 | ** | * |
| 15-20 | ** | * |
| 20-25 | ** | * |
| 25-30 | ** | ns |

| | |
|---|---|
| alpha | 0.05 |
| p value | |
| Time x Refeeding conditions | <0.0001 |
| Time | <0.0001 |
| Refeeding conditions | 0.015 |
| Subject | <0.0001 |
| F (DFn, DFd) | |
| Time x Refeeding conditions | F (24, 1140) = 3.561 |
| Time | F (1.327, 126.1) = 59.54 |
| Refeeding conditions | F (2, 95) = 4.389 |
| Subject | F (95, 1140) = 31.76 |

Fig. 2

Figs. 2A

| Gr5a[-/-]; Gr64a[-/-] | | |
|---|---|---|
| Comparison of Tp between | | p value |
| Fed vs | Starvation | ** |
| | Refed fly food for 10 min | ns |
| | Refed Sucralose for 10 min | *** |
| | Refed Glucose for 10 min | * |
| | Refed Glucose for 1 hr | ns |
| Starvation vs | Refed fly food for 10 min | * |
| | Refed Sucralose for 10 min | ns |
| | Refed Glucose for 10 min | ns |
| | Refed Glucose for 1 hr | ns |

| | |
|---|---|
| p value | <0.0001 |
| alpha | 0.05 |
| Multiple test (ANOVA and Tukey’s post hoc test or Kruskal-Wallis test and Dunn’s test) | Tukey test |
| F value (F (DFn, DFd)) | F (4, 36) = 17.15 |

Fig. 2B

| Gr5a[-/-]; Gr61a[-/-], Gr64a-f[-/-] | | |
|---|---|---|
| Comparison of Tp between | | p value |
| Fed vs | Starvation | **** |
| | Refed fly food for 10 min | ns |
| | Refed Sucralose for 10 min | **** |
| | Refed Glucose for 10 min | *** |
| | Refed Glucose for 1 hr | * |
| Starvation vs | Refed fly food for 10 min | *** |
| | Refed Sucralose for 10 min | ns |
| | Refed Glucose for 10 min | ns |
| | Refed Glucose for 1 hr | ns |

| | |
|---|---|
| p value | P<0.0001 |
| alpha | 0.05 |
| Multiple test (ANOVA and Tukey's post hoc test or Kruskal-Wallis test and Dunn's test) | Tukey test |
| F value (F (DFn, DFd)) | F (5, 32) = 12.07 |

Fig. 2C

| Gr64f-Gal4>uas-Kir | | |
|---|---|---|
| Comparison of Tp between | | p value |
| Fed vs | Starvation | ** |
| | Refed fly food for 10 min | ns |
| | Refed Sucralose for 10 min | * |
| | Refed Glucose for 10 min | ns |
| | Refed Glucose for 1 hr | ns |
| Starvation vs | Refed fly food for 10 min | ** |
| | Refed Sucralose for 10 min | ns |
| | Refed Glucose for 10 min | ns |
| | Refed Glucose for 1 hr | ns |

| | |
|---|---|
| p value | P<0.0001 |
| alpha | 0.05 |
| Multiple test (ANOVA and Tukey's post hoc test or Kruskal-Wallis test and Dunn's test) | Tukey test |
| F value (F (DFn, DFd)) | F (6, 58) = 6.277 |

Fig. 2D

| Gr64fGal4/+ |
|---|

| Comparison of Tp between | | p value |
|---|---|---|
| Fed vs | Starvation | *** |
| | Refed fly food for 10 min | ns |
| | Refed Sucralose for 10 min | ns |
| | Refed Glucose for 10 min | ns |
| Starvation vs | Refed fly food for 10 min | ** |
| | Refed Sucralose for 10 min | * |
| | Refed Glucose for 10 min | ** |

| p value | P=0.0001 |
|---|---|
| alpha | 0.05 |
| Multiple test (ANOVA and Tukey's post hoc test or Kruskal-Wallis test and Dunn's test) | Tukey test |
| F value (F (DFn, DFd)) | F (4, 23) = 9.184 |

Fig. 2E

| uas-Kir/+ | | |
|---|---|---|
| Comparison of Tp between | | p value |
| Fed vs | Starvation | **** |
| | Refed fly food for 10 min | ns |
| | Refed Sucralose for 10 min | * |
| | Refed Glucose for 10 min | ns |
| Starvation vs | Refed fly food for 10 min | **** |
| | Refed Sucralose for 10 min | ** |
| | Refed Glucose for 10 min | **** |

| p value | P<0.0001 |
|---|---|
| alpha | 0.05 |
| Multiple test (ANOVA and Tukey's post hoc test or Kruskal-Wallis test and Dunn's test) | Tukey test |
| F value (F (DFn, DFd)) | F (4, 29) = 20.32 |

Fig. 2G

| Gr64fGal4>uas-CsChrimson | | |
|---|---|---|
| Comparison of Tp between | | p value |
| Fed vs | Water + Red light | **** |
| | ATR + Red light | * |
| Water + Red light vs | ATR + Red light | * |

| | |
|---|---|
| p value | P<0.0001 |
| alpha | 0.05 |
| Multiple test (ANOVA and Tukey’s post hoc test or Kruskal-Wallis test and Dunn’s test) | Tukey test |
| F value (F (DFn, DFd)) | F (2, 23) = 14.35 |

Fig. 2H

| Gr5aGal4>uas-CsChrimson | | |
|---|---|---|
| Comparison of Tp between | | p value |
| Fed vs | Water + Red light | * |
| | ATR + Red light | * |
| Water + Red light vs | ATR + Red light | ns |

| | |
|---|---|
| p value | P=0.0150 |
| alpha | 0.05 |
| Multiple test (ANOVA and Tukey’s post hoc test or Kruskal-Wallis test and Dunn’s test) | Tukey test |
| F value (F (DFn, DFd)) | F (2, 12) = 6.084 |

Fig. 2I

| Gr64aGal4>uas-CsChrimson | | |
|---|---|---|
| Comparison of Tp between | | p value |
| Fed vs | Water + Red light | **** |
| | ATR + Red light | **** |
| Water + Red light vs | ATR + Red light | ns |

| | |
|---|---|
| p value | P<0.0001 |
| alpha | 0.05 |
| Multiple test (ANOVA and Tukey’s post hoc test or Kruskal-Wallis test and Dunn’s test) | Tukey test |
| F value (F (DFn, DFd)) | F (2, 23) = 21.72 |

Fig. 2J

| uas-CsChrimson/+ | | |
|---|---|---|
| Comparison of Tp between | | p value |
| Fed vs | Water + Red light | *** |

| | | |
|---|---|---|
| | ATR + Red light | * |
| Water + Red light vs | ATR + Red light | ns |

| | |
|---|---|
| p value | P=0.0010 |
| alpha | 0.05 |
| Multiple test (ANOVA and Tukey’s post hoc test or Kruskal-Wallis test and Dunn’s test) | Tukey test |
| F value (F (DFn, DFd)) | F (2, 13) = 12.25 |

Fig. 3

Figs. 3A and S7A

| TrpA1[SH]Gal4>uas-Kir | | |
|---|---|---|
| Comparison of Tp between | | p value |
| Fed vs | Starvation | ** |
| | Refed fly food for 10 min | ns |
| | Refed Sucralose for 10 min | **** |
| | Refed Glucose for 10 min | ns |
| | Refed Glucose for 1 hr | ns |
| Starvation vs | Refed fly food for 10 min | * |
| | Refed Sucralose for 10 min | ns |
| | Refed Glucose for 10 min | ns |
| | Refed Glucose for 1 hr | *** |

| | |
|---|---|
| p value | P<0.0001 |
| alpha | 0.05 |
| Multiple test (ANOVA and Tukey’s post hoc test or Kruskal-Wallis test and Dunn’s test) | Tukey test |
| F value (F (DFn, DFd)) | F (5, 42) = 10.02 |

Figs. 3B and S7B

| R11F02Gal4>uas-Kir | | |
|---|---|---|
| Comparison of Tp between | | p value |
| Fed vs | Starvation | * |
| | Refed fly food for 10 min | ns |
| | Refed Sucralose for 10 min | ns |
| | Refed Glucose for 10 min | ns |
| | Refed Glucose for 1 hr | ns |
| Starvation vs | Refed fly food for 10 min | * |
| | Refed Sucralose for 10 min | ns |

| | | |
|---|---|---|
| | Refed Glucose for 10 min | ns |
| | Refed Glucose for 1 hr | ns |

| | |
|---|---|
| p value | 0.0016 |
| alpha | 0.05 |
| Multiple test (ANOVA and Tukey's post hoc test or Kruskal-Wallis test and Dunn's test) | Dunn's test |
| F value (F (DFn, DFd)) | |

Figs. 3C and S7C

| TrpA1[SH]Gal4/+ | | |
|---|---|---|
| Comparison of Tp between | | p value |
| Fed vs | Starvation | ** |
| | Refed fly food for 10 min | ns |
| | Refed Sucralose for 10 min | ns |
| | Refed Glucose for 10 min | ns |
| Starvation vs | Refed fly food for 10 min | ** |
| | Refed Sucralose for 10 min | * |
| | Refed Glucose for 10 min | *** |

| | |
|---|---|
| p value | P=0.0003 |
| alpha | 0.05 |
| Multiple test (ANOVA and Tukey's post hoc test or Kruskal-Wallis test and Dunn's test) | Tukey test |
| F value (F (DFn, DFd)) | F (4, 31) = 7.442 |

Figs. 3D and S7D

| R11F02Gal4/+ | | |
|---|---|---|
| Comparison of Tp between | | p value |
| Fed vs | Starvation | *** |
| | Refed fly food for 10 min | ns |
| | Refed Sucralose for 10 min | ns |
| | Refed Glucose for 10 min | ns |
| Starvation vs | Refed fly food for 10 min | ** |
| | Refed Sucralose for 10 min | * |
| | Refed Glucose for 10 min | *** |

| | |
|---|---|
| p value | P<0.0001 |

| | |
|---|---|
| alpha | 0.05 |
| Multiple test (ANOVA and Tukey's post hoc test or Kruskal-Wallis test and Dunn's test) | Tukey test |
| F value (F (DFn, DFd)) | F (4, 40) = 8.637 |

Fig. 3F

| TrpA1[SH]Gal4>uas-CsChrimson | | |
|---|---|---|
| Comparison of Tp between | | p value |
| Water + Red light vs | ATR + Red light | * |

| | |
|---|---|
| p value | P=0.0186 |
| alpha | 0.05 |
| t-test or Kolmogorov-Smirnov test | Kolmogorov-Smirnov test |

Fig. 3G

| R11F02Gal4>uas-CsChrimson, tubGal80[ts] | | |
|---|---|---|
| Comparison of Tp between | | p value |
| Water + Red light vs | ATR + Red light | * |

| | |
|---|---|
| p value | P=0.0192 |
| alpha | 0.05 |
| t-test or Kolmogorov-Smirnov test | Unpaired t-test |

Fig. 4

Figs. 4A

| NPF[-/-] | | |
|---|---|---|
| Comparison of Tp between | | p value |
| Fed vs | Starvation | **** |
| | Refed fly food for 10 min | *** |
| | Refed Sucralose for 10 min | **** |
| | Refed Glucose for 10 min | **** |
| | Refed Glucose for 1 hr | ** |
| Starvation vs | Refed fly food for 10 min | * |
| | Refed Sucralose for 10 min | ns |
| | Refed Glucose for 10 min | ns |

| | | |
|---|---|---|
| | Refed Glucose for 1 hr | *** |

| | |
|---|---|
| p value | P<0.0001 |
| alpha | 0.05 |
| Multiple test (ANOVA and Tukey's post hoc test or Kruskal-Wallis test and Dunn's test) | Tukey test |
| F value (F (DFn, DFd)) | F (5, 44) = 14.01 |

Figs. 4B

| sNPF hypo | | |
|---|---|---|
| Comparison of Tp between | | p value |
| Fed vs | Starvation | **** |
| | Refed fly food for 10 min | * |
| | Refed Sucralose for 10 min | **** |
| | Refed Glucose for 10 min | *** |
| | Refed Glucose for 1 hr | *** |
| Starvation vs | Refed fly food for 10 min | ** |
| | Refed Sucralose for 10 min | ns |
| | Refed Glucose for 10 min | ** |
| | Refed Glucose for 1 hr | ** |

| | |
|---|---|
| p value | P<0.0001 |
| alpha | 0.05 |
| Multiple test (ANOVA and Tukey's post hoc test or Kruskal-Wallis test and Dunn's test) | Tuckey test |
| F value (F (DFn, DFd)) | F (5, 39) = 18.35 |

Figs. 4C

| Dh44Gal4/+ | | |
|---|---|---|
| Comparison of Tp between | | p value |
| Fed vs | Starvation | **** |
| | Refed fly food for 10 min | ns |
| | Refed Sucralose for 10 min | ** |
| | Refed Glucose for 10 min | ns |
| Starvation vs | Refed fly food for 10 min | **** |
| | Refed Sucralose for 10 min | **** |
| | Refed Glucose for 10 min | **** |

| | |
|---|---|
| p value | P<0.0001 |
| alpha | 0.05 |
| Multiple test (ANOVA and Tukey’s post hoc test or Kruskal-Wallis test and Dunn’s test) | Tukey test |
| F value (F (DFn, DFd)) | F (4, 25) = 27.23 |

Figs. 4D

| Dh44Gal4>uas-Kir | | |
|---|---|---|
| Comparison of Tp between | | p value |
| Fed vs | Starvation | **** |
| | Refed fly food for 10 min | ns |
| | Refed Sucralose for 10 min | **** |
| | Refed Glucose for 10 min | * |
| Starvation vs | Refed fly food for 10 min | **** |
| | Refed Sucralose for 10 min | ns |
| | Refed Glucose for 10 min | *** |

| | |
|---|---|
| p value | P<0.0001 |
| alpha | 0.05 |
| Multiple test (ANOVA and Tukey’s post hoc test or Kruskal-Wallis test and Dunn’s test) | Tukey test |
| F value (F (DFn, DFd)) | F (4, 38) = 21.93 |

Figs. 4E

| AkhGal4/+ | | |
|---|---|---|
| Comparison of Tp between | | p value |
| Fed vs | Starvation | **** |
| | Refed fly food for 10 min | ns |
| | Refed Sucralose for 10 min | ns |
| | Refed Glucose for 10 min | ns |
| Starvation vs | Refed fly food for 10 min | *** |
| | Refed Sucralose for 10 min | *** |
| | Refed Glucose for 10 min | *** |

| | |
|---|---|
| p value | P<0.0001 |
| alpha | 0.05 |
| Multiple test (ANOVA and Tukey’s post hoc test or Kruskal-Wallis test and Dunn’s test) | Tukey test |
| F value (F (DFn, DFd)) | F (4, 30) = 14.32 |

Figs. 4F

| AkhGal4>uas-Kir | | |
|---|---|---|
| Comparison of Tp between | | p value |
| Fed vs | Starvation | **** |
| | Refed fly food for 10 min | ns |
| | Refed Sucralose for 10 min | *** |
| | Refed Glucose for 10 min | ns |
| | Refed Glucose for 1 hr | ns |
| Starvation vs | Refed fly food for 10 min | **** |
| | Refed Sucralose for 10 min | ns |
| | Refed Glucose for 10 min | **** |
| | Refed Glucose for 1 hr | **** |

| | |
|---|---|
| p value | P<0.0001 |
| alpha | 0.05 |
| Multiple test (ANOVA and Tukey’s post hoc test or Kruskal-Wallis test and Dunn’s test) | Tukey test |
| F value (F (DFn, DFd)) | F (5, 35) = 16.47 |

Figs. 4G

| ilp6 LOF | | |
|---|---|---|
| Comparison of Tp between | | p value |
| Fed vs | Starvation | **** |
| | Refed fly food for 10 min | ns |
| | Refed Sucralose for 10 min | **** |
| | Refed Glucose for 10 min | ns |
| | Refed Glucose for 1 hr | ns |
| Starvation vs | Refed fly food for 10 min | **** |
| | Refed Sucralose for 10 min | ns |
| | Refed Glucose for 10 min | **** |
| | Refed Glucose for 1 hr | **** |

| | |
|---|---|
| p value | P<0.0001 |
| alpha | 0.05 |
| Multiple test (ANOVA and Tukey’s post hoc test or Kruskal-Wallis test and Dunn’s test) | Tukey test |
| F value (F (DFn, DFd)) | F (5, 48) = 23.13 |

Figs. 4H

| Upd3Δ | | |
|---|---|---|
| Comparison of Tp between | | p value |
| Fed vs | Starvation | **** |
| | Refed fly food for 10 min | ns |
| | Refed Sucralose for 10 min | **** |
| | Refed Glucose for 10 min | ns |
| | Refed Glucose for 1 hr | ns |
| Starvation vs | Refed fly food for 10 min | ns |
| | Refed Sucralose for 10 min | ns |
| | Refed Glucose for 10 min | * |
| | Refed Glucose for 1 hr | ** |

| | |
|---|---|
| p value | P<0.0001 |
| alpha | 0.05 |
| Multiple test (ANOVA and Tukey's post hoc test or Kruskal-Wallis test and Dunn's test) | Tukey test |
| F value (F (DFn, DFd)) | F (5, 37) = 14.92 |

Figs. 4I

| Upd2Δ | | |
|---|---|---|
| Comparison of Tp between | | p value |
| Fed vs | Starvation | ** |
| | Refed fly food for 10 min | ns |
| | Refed Sucralose for 10 min | **** |
| | Refed Glucose for 10 min | ns |
| | Refed Glucose for 1 hr | ns |
| Starvation vs | Refed fly food for 10 min | ns |
| | Refed Sucralose for 10 min | ns |
| | Refed Glucose for 10 min | ns |
| | Refed Glucose for 1 hr | ns |

| | |
|---|---|
| p value | P=0.0002 |
| alpha | 0.05 |
| Multiple test (ANOVA and Tukey's post hoc test or Kruskal-Wallis test and Dunn's test) | Tukey test |
| F value (F (DFn, DFd)) | F (5, 43) = 6.356 |

Fig. 5
Fig. 5A
w[1118], LD

| w[1118], LD, ZT1-3 | | |
|---|---|---|
| Comparison of Tp between | | p value |
| Fed vs | Starvation | *** |
| | Refed Sucralose for 10 min | ** |
| | Refed Glucose for 10 min | NS |
| | Refed Glucose for 1 hr | NS |
| Starvation vs | Refed Sucralose for 10 min | NS |
| | Refed Glucose for 10 min | **** |
| | Refed Glucose for 1 hr | **** |

| | |
|---|---|
| p value | P<0.0001 |
| alpha | 0.05 |
| Multiple test (ANOVA and Tukey's post hoc test or Kruskal-Wallis test and Dunn's test) | Tukey test |
| F value (F (DFn, DFd)) | F (4, 26) = 14.24 |

| w[1118], LD, ZT4-6 | | |
|---|---|---|
| Comparison of Tp between | | p value |
| Fed vs | Starvation | **** |
| | Refed Sucralose for 10 min | NS |
| | Refed Glucose for 10 min | NS |
| | Refed Glucose for 1 hr | NS |
| Starvation vs | Refed Sucralose for 10 min | ** |
| | Refed Glucose for 10 min | *** |
| | Refed Glucose for 1 hr | *** |

| | |
|---|---|
| p value | P<0.0001 |
| alpha | 0.05 |
| Multiple test (ANOVA and Tukey's post hoc test or Kruskal-Wallis test and Dunn's test) | Tukey test |
| F value (F (DFn, DFd)) | F (4, 36) = 9.693 |

| w[1118], LD, ZT7-9 | | |
|---|---|---|
| Comparison of Tp between | | p value |
| Fed vs | Starvation | ** |

| | | |
|---|---|---|
| | Refed Sucralose for 10 min | NS |
| | Refed Glucose for 10 min | NS |
| | Refed Glucose for 1 hr | NS |
| Starvation vs | Refed Sucralose for 10 min | NS |
| | Refed Glucose for 10 min | NS |
| | Refed Glucose for 1 hr | * |

| | |
|---|---|
| p value | P=0.0054 |
| alpha | 0.05 |
| Multiple test (ANOVA and Tukey's post hoc test or Kruskal-Wallis test and Dunn's test) | Tukey test |
| F value (F (DFn, DFd)) | F (4, 25) = 4.768 |

| w[1118], LD, ZT10-12 | | |
|---|---|---|
| Comparison of Tp between | | p value |
| Fed vs | Starvation | **** |
| | Refed Sucralose for 10 min | * |
| | Refed Glucose for 10 min | NS |
| | Refed Glucose for 1 hr | NS |
| Starvation vs | Refed Sucralose for 10 min | **** |
| | Refed Glucose for 10 min | **** |
| | Refed Glucose for 1 hr | **** |

| | |
|---|---|
| p value | P=0.0054 |
| alpha | 0.05 |
| Multiple test (ANOVA and Tukey's post hoc test or Kruskal-Wallis test and Dunn's test) | Tukey test |
| F value (F (DFn, DFd)) | F (4, 25) = 4.768 |

Fig. 5B
y[1]w[1], LD

| y[1]w[1], LD, ZT1-3 | | |
|---|---|---|
| Comparison of Tp between | | p value |
| Fed vs | Starvation | **** |
| | Refed Sucralose for 10 min | ns |
| | Refed Glucose for 10 min | ns |
| Starvation vs | Refed Sucralose for 10 min | *** |
| | Refed Glucose for 10 min | ** |

| | |
|---|---|
| p value | P<0.0001 |
| alpha | 0.05 |
| Multiple test (ANOVA and Tukey's post hoc test or Kruskal-Wallis test and Dunn's test) | Tukey test |
| F value (F (DFn, DFd)) | F (3, 32) = 12.04 |

| y[1]w[1], LD, ZT4-6 | | |
|---|---|---|
| Comparison of Tp between | | p value |
| Fed vs | Starvation | ** |
| | Refed Sucralose for 10 min | ns |
| | Refed Glucose for 10 min | ns |
| Starvation vs | Refed Sucralose for 10 min | * |
| | Refed Glucose for 10 min | ** |
| p value | | P<0.0001 |
| alpha | | 0.05 |
| Multiple test (ANOVA and Tukey's post hoc test or Kruskal-Wallis test and Dunn's test) | | Dunn's test |
| F value (F (DFn, DFd)) | | |

| y[1]w[1], LD, ZT7-9 | | |
|---|---|---|
| Comparison of Tp between | | p value |
| Fed vs | Starvation | **** |
| | Refed Sucralose for 10 min | NS |
| | Refed Glucose for 10 min | NS |
| Starvation vs | Refed Sucralose for 10 min | *** |
| | Refed Glucose for 10 min | **** |

| | |
|---|---|
| p value | P<0.0001 |
| alpha | 0.05 |
| Multiple test (ANOVA and Tukey's post hoc test or Kruskal-Wallis test and Dunn's test) | Tukey test |
| F value (F (DFn, DFd)) | F (3, 32) = 23.95 |

| y[1]w[1], LD, ZT10-12 | | |
|---|---|---|
| Comparison of Tp between | | p value |
| Fed vs | Starvation | **** |
| | Refed Sucralose for 10 min | NS |
| | Refed Glucose for 10 min | NS |
| Starvation vs | Refed Sucralose for 10 min | **** |

| | | |
|---|---|---|
| | Refed Glucose for 10 min | **** |

| | |
|---|---|
| p value | P=0.0001 |
| alpha | 0.05 |
| Multiple test (ANOVA and Tukey's post hoc test or Kruskal-Wallis test and Dunn's test) | Dunn's test |
| F value (F (DFn, DFd)) | |

Fig. 5C
per[01], LD

| per[01], LD, ZT1-3 | | |
|---|---|---|
| Comparison of Tp between | | p value |
| Fed vs | Starvation | ** |
| | Refed Sucralose for 10 min | **** |
| | Refed Glucose for 10 min | ns |
| Starvation vs | Refed Sucralose for 10 min | ns |
| | Refed Glucose for 10 min | **** |

| | |
|---|---|
| p value | P<0.0001 |
| alpha | 0.05 |
| Multiple test (ANOVA and Tukey's post hoc test or Kruskal-Wallis test and Dunn's test) | Tukey test |
| F value (F (DFn, DFd)) | F (3, 23) = 28.56 |

| per[01], LD, ZT4-6 | | |
|---|---|---|
| Comparison of Tp between | | p value |
| Fed vs | Starvation | * |
| | Refed Sucralose for 10 min | ** |
| | Refed Glucose for 10 min | ns |
| Starvation vs | Refed Sucralose for 10 min | ns |
| | Refed Glucose for 10 min | ** |

| | |
|---|---|
| p value | P=0.0003 |
| alpha | 0.05 |
| Multiple test (ANOVA and Tukey's post hoc test or Kruskal-Wallis test and Dunn's test) | Tukey test |
| F value (F (DFn, DFd)) | F (3, 22) = 9.425 |

| per[01], LD, ZT7-9 | | |
|---|---|---|
| Comparison of Tp between | | p value |
| Fed vs | Starvation | ** |
| | Refed Sucralose for 10 min | * |
| | Refed Glucose for 10 min | ns |
| Starvation vs | Refed Sucralose for 10 min | ns |
| | Refed Glucose for 10 min | * |

| | |
|---|---|
| p value | P=0.0014 |
| alpha | 0.05 |
| Multiple test (ANOVA and Tukey's post hoc test or Kruskal-Wallis test and Dunn's test) | Dunn's test |
| F value (F (DFn, DFd)) | |

| per[01], LD, ZT10-12 | | |
|---|---|---|
| Comparison of Tp between | | p value |
| Fed vs | Starvation | * |
| | Refed Sucralose for 10 min | *** |
| | Refed Glucose for 10 min | ns |
| Starvation vs | Refed Sucralose for 10 min | ns |
| | Refed Glucose for 10 min | * |

| | |
|---|---|
| p value | P<0.0001 |
| alpha | 0.05 |
| Multiple test (ANOVA and Tukey's post hoc test or Kruskal-Wallis test and Dunn's test) | Tukey test |
| F value (F (DFn, DFd)) | F (3, 17) = 13.36 |

Fig. 5D
tim[01], LD

| tim[01], LD, ZT1-3 |
|---|

| Comparison of Tp between | | p value |
|---|---|---|
| Fed vs | Starvation | **** |
| | Refed Sucralose for 10 min | **** |
| | Refed Glucose for 10 min | ns |
| Starvation vs | Refed Sucralose for 10 min | ns |
| | Refed Glucose for 10 min | **** |

| | |
|---|---|
| p value | P<0.0001 |
| alpha | 0.05 |
| Multiple test (ANOVA and Tukey's post hoc test or Kruskal-Wallis test and Dunn's test) | Tukey test |
| F value (F (DFn, DFd)) | F (3, 18) = 34.89 |

| tim[01], LD, ZT4-6 | | |
|---|---|---|
| Comparison of Tp between | | p value |
| Fed vs | Starvation | **** |
| | Refed Sucralose for 10 min | **** |
| | Refed Glucose for 10 min | ns |
| Starvation vs | Refed Sucralose for 10 min | ns |
| | Refed Glucose for 10 min | **** |

| | |
|---|---|
| p value | P<0.0001 |
| alpha | 0.05 |
| Multiple test (ANOVA and Tukey's post hoc test or Kruskal-Wallis test and Dunn's test) | Tukey test |
| F value (F (DFn, DFd)) | F (3, 32) = 44.40 |

| tim[01], LD, ZT7-9 | | |
|---|---|---|
| Comparison of Tp between | | p value |
| Fed vs | Starvation | **** |
| | Refed Sucralose for 10 min | **** |
| | Refed Glucose for 10 min | ns |
| Starvation vs | Refed Sucralose for 10 min | ns |
| | Refed Glucose for 10 min | **** |

| | |
|---|---|
| p value | P<0.0001 |
| alpha | 0.05 |

| | |
|---|---|
| Multiple test (ANOVA and Tukey's post hoc test or Kruskal-Wallis test and Dunn's test) | Tukey test |
| F value (F (DFn, DFd)) | F (3, 39) = 28.77 |

| tim[01], LD, ZT10-12 | | |
|---|---|---|
| Comparison of Tp between | | p value |
| Fed vs | Starvation | *** |
| | Refed Sucralose for 10 min | ** |
| | Refed Glucose for 10 min | ns |
| Starvation vs | Refed Sucralose for 10 min | ns |
| | Refed Glucose for 10 min | * |

| | |
|---|---|
| p value | P=0.0002 |
| alpha | 0.05 |
| Multiple test (ANOVA and Tukey's post hoc test or Kruskal-Wallis test and Dunn's test) | Dunn's test |
| F value (F (DFn, DFd)) | |

Fig. S3

| w[1118] | | |
|---|---|---|
| Comparison of Tp between | | p value |
| Fed vs | Starvation for 1 ON (STV1ON) | **** |
| | STV1ON+Refed Sucralose for 10 min | *** |
| | STV1ON+Refed Glucose for 10 min | ns |
| STV1ON vs | STV1ON+Refed Sucralose for 10 min | ** |
| | STV1ON+Refed Glucose for 10 min | **** |
| Fed vs | Starvation for 1.5 ON (STV1.5ON) | **** |
| | STV1.5ON+Refed Sucralose for 10 min | ** |
| | STV1.5ON+Refed Glucose for 10 min | ns |
| STV1.5ON vs | STV1.5ON+Refed Sucralose for 10 min | ** |
| | STV1.5ON+Refed Glucose for 10 min | **** |

| | |
|---|---|
| p value | P<0.0001 |
| alpha | 0.05 |
| Multiple test (ANOVA and Tukey's post hoc test or Kruskal-Wallis test and Dunn's test) | Tukey test |
| F value (F (DFn, DFd)) | F (6, 42)=23.02 |

Fig. S4

| *Orco*[1] | | |
|---|---|---|
| Comparison of Tp between | | p value |
| Fed vs | Starvation | **** |
| | Refed fly food for 10 min | *** |
| | Refed Sucralose for 10 min | **** |
| | Refed Glucose for 10 min | *** |
| | Refed Glucose for 1 hr | ns |
| Starvation vs | Refed fly food for 10 min | * |
| | Refed Sucralose for 10 min | ns |
| | Refed Glucose for 10 min | * |
| | Refed Glucose for 1 hr | **** |

| | |
|---|---|
| p value | P<0.0001 |
| alpha | 0.05 |
| Multiple test (One-way Anova and Tukey's test or Kraskal-Wallis test and Dunn's test) | Tuckey's test |
| F value (F (DFn, DFd)) | F (5, 49) = 18.90 |

Fig. S5
w[1118], DD

| w[1118], DD, CT1-3 | | |
|---|---|---|
| Comparison of Tp between | | p value |
| Fed vs | Starvation | ** |
| | Refed Sucralose for 10 min | ** |
| | Refed Glucose for 10 min | NS |
| Starvation vs | Refed Sucralose for 10 min | NS |
| | Refed Glucose for 10 min | ** |

| | |
|---|---|
| p value | P<0.0001 |
| alpha | 0.05 |
| Multiple test (One-way Anova and Tuckey's test or Kraskal-Wallis test and Dunn’s test) | Dunn's test |
| F value (F (DFn, DFd)) | |

| w[1118], DD, CT4-6 | | |
|---|---|---|
| Comparison of Tp between | | p value |
| Fed vs | Starvation | **** |
| | Refed Sucralose for 10 min | **** |
| | Refed Glucose for 10 min | ns |
| Starvation vs | Refed Sucralose for 10 min | ns |
| | Refed Glucose for 10 min | **** |

| | |
|---|---|
| p value | P<0.0001 |
| alpha | 0.05 |
| Multiple test (One-way Anova and Tuckey's test or Kraskal-Wallis test and Dunn’s test) | Tuckey's test |
| F value (F (DFn, DFd)) | F (3, 40) = 21.31 |

| w[1118], DD, CT7-9 | | |
|---|---|---|
| Comparison of Tp between | | p value |
| Fed vs | Starvation | **** |
| | Refed Sucralose for 10 min | **** |
| | Refed Glucose for 10 min | ns |
| Starvation vs | Refed Sucralose for 10 min | ns |
| | Refed Glucose for 10 min | **** |

| | |
|---|---|
| p value | P<0.0001 |
| alpha | 0.05 |
| Multiple test (One-way Anova and Tuckey's test or Kraskal-Wallis test and Dunn’s test) | Tuckey's test |
| F value (F (DFn, DFd)) | F (3, 37) = 25.78 |

| w[1118], DD, CT10-12 | | |
|---|---|---|
| Comparison of Tp between | | p value |

| | | |
|---|---|---|
| Fed vs | Starvation | **** |
| | Refed Sucralose for 10 min | *** |
| | Refed Glucose for 10 min | ns |
| Starvation vs | Refed Sucralose for 10 min | ns |
| | Refed Glucose for 10 min | ** |

| | |
|---|---|
| p value | <0.0001 |
| alpha | 0.05 |
| Multiple test (One-way Anova and Tuckey's test or Kraskal-Wallis test and Dunn's test) | Dunn's test |
| F value (F (DFn, DFd)) | |

Fig. S6

Fig. S6A

| *yw, Cumulative # of licking* | | |
|---|---|---|
| Time | Starvation vs Refed Sucralose | Starvation vs Refed Glucose |
| 0-5 | ns | ns |
| 5-10 | ns | ns |
| 10-15 | ns | ns |
| 15-20 | ns | ns |
| 20-25 | ns | ns |
| 25-30 | ns | ns |

| | |
|---|---|
| alpha | 0.05 |
| p value | |
| Time x Refeeding conditions) | 0.9999 |
| Time | <0.0001 |
| Refeeding conditions | 0.8321 |
| Subject | <0.0001 |
| F (DFn, DFd) | |
| Time x Refeeding conditions | F (24, 1704) = 0.2587 |
| Time | F (1.189, 168.8) = 102.8 |
| Refeeding conditions | F (2, 142) = 0.1841 |
| Subject | F (142, 1704) = 27.38 |

Fig. S6B

| *$per^{01}$, Cumulative # of licking* | | |
|---|---|---|
| Time | Starvation vs Refed Sucralose | Starvation vs Refed Glucose |

| | | |
|---|---|---|
| 0-5 | *** | ns |
| 5-10 | ** | ns |
| 10-15 | ** | ns |
| 15-20 | *** | ns |
| 20-25 | *** | ns |
| 25-30 | *** | ns |

| | |
|---|---|
| alpha | 0.05 |
| p value | |
| Time x Refeeding conditions) | <0.0001 |
| Time | <0.0001 |
| Refeeding conditions | 0.0018 |
| Subject | <0.0001 |
| F (DFn, DFd) | |
| Time x Refeeding conditions | F (24, 1476) = 6.720 |
| Time | F (1.137, 139.9) = 73.68 |
| Refeeding conditions | F (2, 123) = 6.686 |
| Subject | F (123, 1476) = 32.81 |

Fig. S6C

| *$tim^{01}$, Cumulative # of licking* | | |
|---|---|---|
| Time | Starvation vs Refed Sucralose | Starvation vs Refed Glucose |
| 0-5 | ns | * |
| 5-10 | ns | ** |
| 10-15 | ns | * |
| 15-20 | ns | * |
| 20-25 | ns | * |
| 25-30 | ns | ns |

| | |
|---|---|
| alpha | 0.05 |
| p value | |
| Time x Refeeding conditions) | 0.5253 |
| Time | <0.0001 |
| Refeeding conditions | 0.211 |
| Subject | <0.0001 |
| F (DFn, DFd) | |
| Time x Refeeding conditions | F (24, 1668) = 0.9547 |
| Time | F (1.182, 164.2) = 190.0 |
| Refeeding conditions | F (2, 139) = 1.574 |
| Subject | F (139, 1668) = 27.06 |
